# Structural basis for lipid transport at membrane contact sites by the Ist2/Osh6 complex

**DOI:** 10.1101/2025.02.18.638816

**Authors:** Melanie Arndt, Angela Schweri, Raimund Dutzler

## Abstract

Membrane contact sites (MCS) are hubs for inter-organellar lipid transport within eukaryotic cells. As one of the principal tethers bridging the ER and the plasma membrane in *Saccharomyces cerevisiae*, the protein Ist2 plays a major role during lipid transport between both compartments. Here, we present a comprehensive investigation elucidating structural and mechanistic properties of Ist2 and its interaction with the soluble lipid transport protein Osh6. The ER-embedded transmembrane domain of Ist2 is homologous to the TMEM16 family and acts as constitutively active lipid scramblase. The extended C-terminus binds to the plasma membrane and the PS/PI_4_P exchanger Osh6. Through cellular growth assays, biochemical and structural studies, we have elucidated the interaction between both proteins and provide evidence that Osh6 remains associated with Ist2 during lipid shuttling between membranes. These results highlight the role of the Ist2/Osh6 complex in lipid trafficking and offer new insights into the relevance of scramblases at MCSs.

---

The lipid composition of membranes surrounding cellular compartments is highly heterogeneous, resulting from a combination of localized biosynthesis and complex transport processes^1–3^. The negatively charged phosphatidylserine (PS), for example, is synthesized in the endoplasmic reticulum (ER), while it is accumulated in the plasma membrane (PM)^4–6^. In eukaryotic cells, the transport of PS is largely mediated by members of the oxysterol-binding protein related protein family (ORP), which contain a specific lipid binding site that allows the exchange of lipids at regions where membranes are in close apposition^7–11^. In the yeast *Saccharomyces cerevisiae*, the transport of PS to the plasma membrane is mediated by the ORP homologues Osh6 and Osh7, which accumulate PS on the plasma membrane in exchange for the phosphoinositide PI_4_P^9,11^. PI_4_P is transferred from the PM to the ER, where its dephosphorylation by the phosphatase Sac1 helps to maintain a concentration gradient that drives transport^12^. This process takes place at the cortical ER, which is abundant in *S. cerevisiae* and stabilized by different tethering proteins that keep both membranes at distances ranging between 10-30 nm as revealed from electron microscopy data^13–15^. Since Osh6 and Osh7 themselves lack domains localizing them to the cortical ER, they bind to Ist2, which is one of the primary tethers found to form ER-PM contact sites in this unicellular organism^16–19^.

The 946-residues long Ist2 is a member of the TMEM16 family, encompassing ion channels and lipid scramblases in eukaryotic cells that are regulated by intracellular calcium^20–22^. Ist2 distinguishes itself from other family members by its modular organization, consisting of a membrane-inserted N-terminal half and an extended C-terminal cytoplasmic region of comparable length. The sequence of the N-terminal part resembles other fungal TMEM16 scramblases and the mammalian paralogs TMEM16H and K, although without the residues of the Ca^2+^ binding site, which are strongly conserved among other family members^23^. In contrast, the following region is predicted to be unstructured and to bridge the ER with the plasma membrane by binding to the inner leaflet of the latter via a cluster of positively charged residues at the C-terminal end^19^. The cytoplasmic region of Ist2 was shown to localize Osh6 and 7 to membrane contact sites (MCS) via binding to a stretch in the center of the elongated tail, which is conserved within the family of *Saccharomycetaceae*^16,17^. In the following, we will focus on Osh6, but expect our findings also to extend towards the highly homologous Osh7. While the coarse region interacting with Osh6 has been identified, the details of the Ist2-Osh6 interaction and its role in non-vesicular lipid transport has remained elusive. Open questions concern the function of Ist2 as lipid scramblase, the structural basis of its interaction with Osh6 and whether the two proteins remain associated during lipid shuttling. To address these questions, we have engaged in extended investigations using a combination of biochemical, structural and cellular approaches to understand the intricate interplay between Ist2 and Osh6 at MCS.

## Results

### Structural Features of the Ist2/Osh6 complex

In our study, we aimed for a detailed comprehension of the Ist2/Osh6 system for lipid transport. To characterize the complex *in vitro*, we have separately over-expressed full-length Ist2 and Osh6 in *S. cerevisiae*. After their purification and reconstitution, we found both proteins to co-elute as complex on size exclusion chromatography (SEC, Fig. 1a). A structural characterization of this sample by cryo-EM provided distinct reconstructions that can be attributed to the transmembrane domain of Ist2 and bound Osh6, respectively, in accordance with the predicted modular organization of Ist2 consisting of a folded N-terminal and a largely unstructured C-terminal halve (Fig. 1b, Extended Data Fig. 1, Extended Data Table 1, Supplementary Fig. 1). The membrane-inserted domain of Ist2 assembles as dimer showing the characteristic architecture of a TMEM16 protein^23^ (Fig. 1c, Extended Data Fig. 1a). In the case of the bound Osh6, the small size of the protein has compromised particle alignment and consequently limited the obtained resolution (Fig. 1d, Extended Data Fig. 1a). In this dataset, we also found 2D classes containing multiple particles at short distances, which could account for potential interactions between lipid transfer proteins (LTP) and Ist2 (Extended Data Fig. 1a).

**Figure 1.**
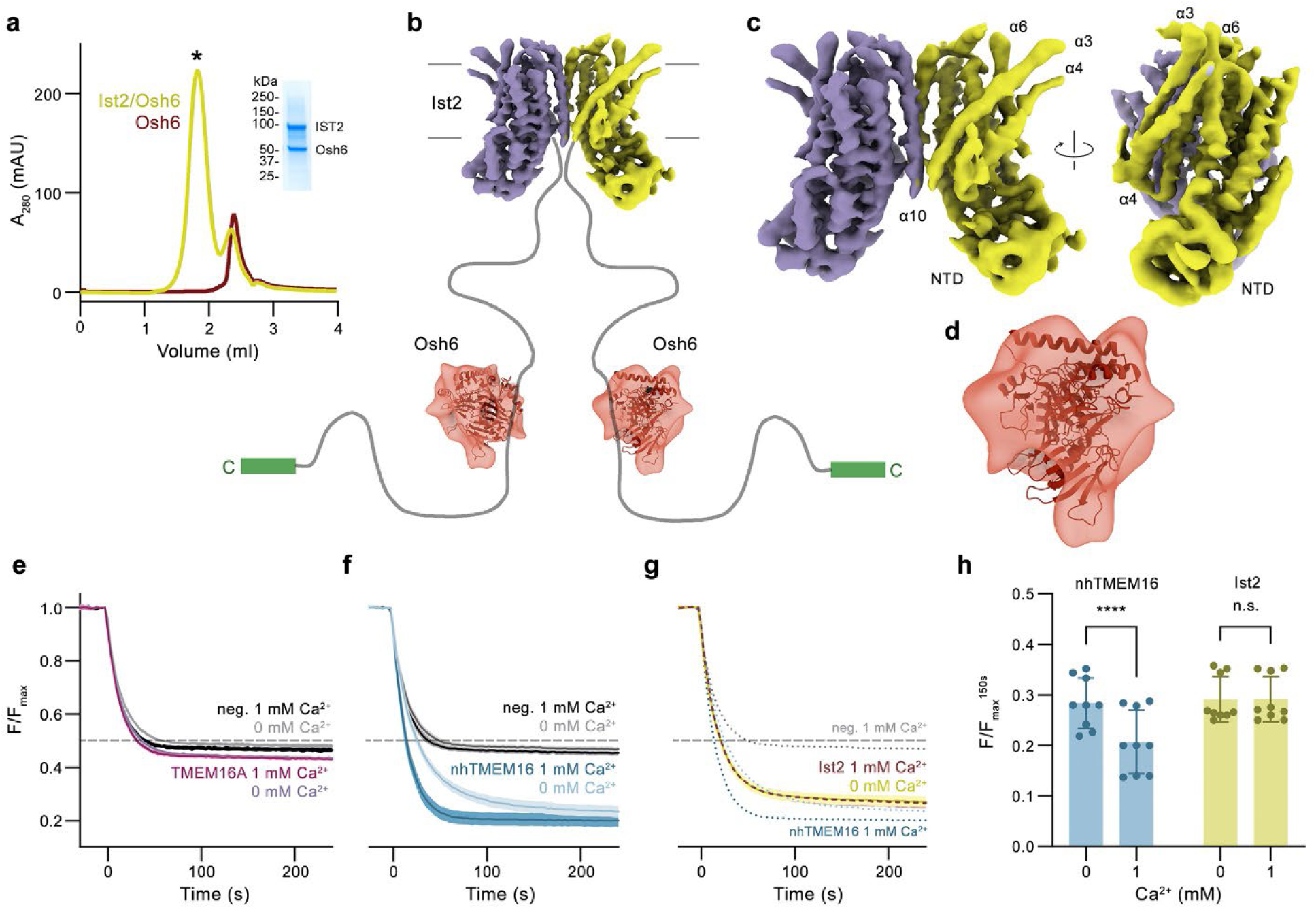
Structural and functional properties of the Ist2/Osh6 complex. **a**, SEC profile of a sample containing Ist2 and Osh6 at a 1.5 molar excess (yellow). Peak fraction (*) contains the complex of both proteins as shown by SDS PAGE (inset right). Red trace represents free Osh6. **b**, Scheme composed of the cryo-EM density of Ist2, and Osh6 as observed in their complex with flexible linker that is not defined in the structure indicated. The C-terminal parts interacting with the PM are shown in green, the membrane boundaries are indicated. **c**, Cryo-EM density of the N-terminal membrane-inserted part of the Ist2 dimer at 4.9 Å. **d**, Cryo-EM density of Osh6 at 10.5 Å with a placed ribbon representation of its structure (PDBID: 4B2Z) shown for comparison. **e-h**, Lipid transport properties of the N-terminal construct Ist2^716^ assayed in liposomes as the irreversible bleaching of fluorescent lipids facing the outside. **e**, Absence of lipid transport activity in mTMEM16A-containing proteoliposomes and liposomes not containing any protein (neg.) shown as control. Data show averages of three measurements obtained from one reconstitution, errors are s.e.m. **f**, Ca^2+^-dependent scrambling in proteoliposomes containing nhTMEM16 and liposomes not containing any protein (neg.) shown as control. **g**, Ca^2+^-independent scrambling in proteoliposomes containing Ist2^716^. Traces of empty liposomes and nhTMEM16 (displayed in b) are shown as dotted lines for comparison. **a**, **b**, Data show averages of nine measurements obtained from three independent reconstitutions, errors are s.e.m. **a**-**c** Traces depict fluorescence decrease of tail-labeled NBD-PE lipids after addition of dithionite (t=0) in absence and presence of 1mM Ca^2+^. Dashed grey line indicates 0.5 threshold. **h**, Comparison of the fluorescence 150 s after addition of dithionite in proteoliposomes containing either nhTMEM16 or Ist2^716^. Bars show average of nine experiments (displayed as dots), errors are s.e.m.. The fluorescence difference in presence of Ca^2+^ is significant in case of nhTMEM16 (p<0.001) but not Ist2^716^ (p>0.9999) as calculated by ANOVA.

### The N-terminal domain of Ist2 forms a constitutively active lipid scramblase

Next, we were interested whether the N-terminal part of Ist2 forms a functionally autonomous unit that has retained its capability to mediate lipid transfer between both leaflets of the membrane, in spite of a previous publication refuting scrambling activity of the protein^24^. We thus prepared a construct containing the first 716 amino acids (Ist2^716^), which lacks the bulk of the C-terminal region including the proposed Osh6 binding sequence (Extended Data Fig. 2a, Supplementary Fig. 1a). To investigate its lipid transport capability, we have reconstituted the purified Ist2^716^ into liposomes containing trace amounts of fluorescent lipids (Extended Data Fig. 2b). Scrambling was assayed by following the time dependent decrease of the fluorescence upon bleaching of labeled lipids residing in the outer leaflet of the bilayer^24^ (Extended Data Fig. 2c). In case of empty liposomes or proteoliposomes containing TMEM16A, which is unable to transport lipids, the fluorescence decays to about 50% of its initial value (Fig. 1e). In contrast, in the presence of an active scramblase, the fluorescence plateau is further decreased as a consequence of the lipid transfer between both leaflets (Fig. 1f). For the fungal scramblase nhTMEM16, we observe a slow fluorescence decay to 15-30% of its initial value in the absence of Ca^2+^, and faster kinetics upon Ca^2+^addition, reflecting the increased protein activity upon ligand binding (Fig. 1f, Extended Data Fig. 2d-f)^23^. Reconstituted Ist2^716^ already displays pronounced scrambling activity in the absence of Ca^2+^, which is not further enhanced upon Ca^2+^ addition, which distinguishes the protein as constitutively active lipid scramblase (Fig. 1g, h, Extended Data Fig. 2b, d-f). This feature aligns with the absence of the conserved Ca^2+^ binding site in homologues from *Saccharomycetaceae*, which are all supposed to participate in similar processes at MCSs (Supplementary Fig. 1b).

### Structure of the Ist2 Scramblase unit

After identifying the membrane-inserted part of Ist2 as constitutively active scramblase, we were interested in how this functional feature would be manifested in its architecture. Since we did not succeed in obtaining a high-resolution structure of the full-length protein, we used the well-behaved Ist2^716^ construct for cryo-EM sample preparation, for which we obtained reconstructions in the detergent GDN at a global resolution of 2.84 Å and in a small-sized lipid nanodiscs (assembled with the lipid scaffolding protein 1E3D1) at 3.35 and 3.8 Å, respectively (Fig. 2a, Extended Data Figs. 2g, h, 3 and 4, Extended Data Table 1). As observed for the transmembrane domain of the full-length protein, Ist2^716^ is a homodimer that shares the general features of known structures of TMEM16 family members^23,25–28^ (Fig. 2a, b). Each subunit consists of a cytosolic N-terminal domain (NTD) that is followed by a unit comprising 10 membrane-spanning segments and a disordered C-terminus encompassing residues 596-716 (Fig. 2b, c, Supplementary Fig. 1a). Although the NTD of Ist2 resembles the equivalent region of nhTMEM16, this unit is considerably smaller and restricted to the core ferredoxin fold consisting of four-stranded antiparallel β-sheet and two interspersed α-helices (Fig. 2c, Extended Data Fig. 5a). This cytoplasmic domain tightly interacts with intracellular helices and loops at the periphery of Ist2 remote from the dimer interface to stabilize the observed structure in this region (Fig. 2c, Extended Data Fig. 5b). The dimer interface burying 1516 Å^2^ of the combined molecular surface involves contacts between the symmetry-related helices α10 located in the outer- and between α10 and α3 at the inner leaflet of the bilayer (Extended Data Fig. 5c). The low-resolution density of the detergent micelle surrounding the membrane-embedded part of the protein shows a pronounced distortion that was previously described as hallmark of family members acting as lipid scramblases^26,29,30^ (Fig. 2d). Both boundaries of the distorted micelle delineate a trail of lipid-like densities surrounding the TM-helices that is still visible at high map contour (Fig. 2d, Extended Data Fig. 5d). A similar distortion is also observed in two distinct reconstructions from a dataset of Ist2^716^ in lipid nanodiscs (Extended Data Fig. 4 and 5e), which further underlines the role of the protein in destabilizing its environment. Bound detergent molecules and lipids follow the arrangement of transmembrane helices, resulting in an apparent thinning of the bilayer at both entrances of the subunit cavity, closely resembling previously described features observed in the fungal scramblases afTMEM16 and nhTMEM16^26,29,30^ (Fig. 2d, Extended Data Fig. 5d, e). Similarly, the part of the protein defining the subunit cavity, which is comprised of α-helices 3-8, displays a striking resemblance to the Ca^2+^-bound active conformations of nhTMEM16, afTMEM16 and human TMEM16K^23,27,29^ (Fig. 2e, Extended Data Fig. 5f). By forming a continuous polar furrow exposed to the core of the membrane, which is confined by α-helices 4 and 6, this unit is of appropriate width to accommodate a lipid headgroup, presumably lowering the barrier for lipid flip-flop (Fig. 2e, Extended Data Fig. 5g). Despite the described similarities, Ist2 contains several unique structural hallmarks, which presumably stabilize its conformation to impede a collapse of the polar furrow as observed in other fungal family members in absence of Ca^2+^ (ref. ^26,27,29^). In nhTMEM16 and its orthologs, the open conformation is stabilized by Ca^2+^ binding to an extended site located within the inner membrane leaflet that is strongly conserved within the TMEM16 family^23^. This site contains five acidic residues on α-helices 6-8 that coordinate two Ca^2+^ ions to rigidify the intracellular part of α-helix 6 in its interaction with α8 (Fig. 2f). In Ist2, three of the conserved acidic residues located on α7 and 8 are substituted. In particular, the swap of a glutamate (Glu 506 in nhTMEM16) with a lysine (Lys 443 in Ist2) is striking, as it introduces a positively charged sidechain that replaces Ca^2+^ by forming a bifurcated salt bridge with the conserved residues Glu 374 on α6 and Asp 476 on α8, (Fig. 2f). Additionally, the observed conformation of α6, whose lower part shows a stronger bending towards the membrane plane compared to other homologs of known structure, is stabilized by a successive short cytoplasmic helix (α6_i_) in a region that is disordered in other family members (Fig. 2g).

**Figure 2.**
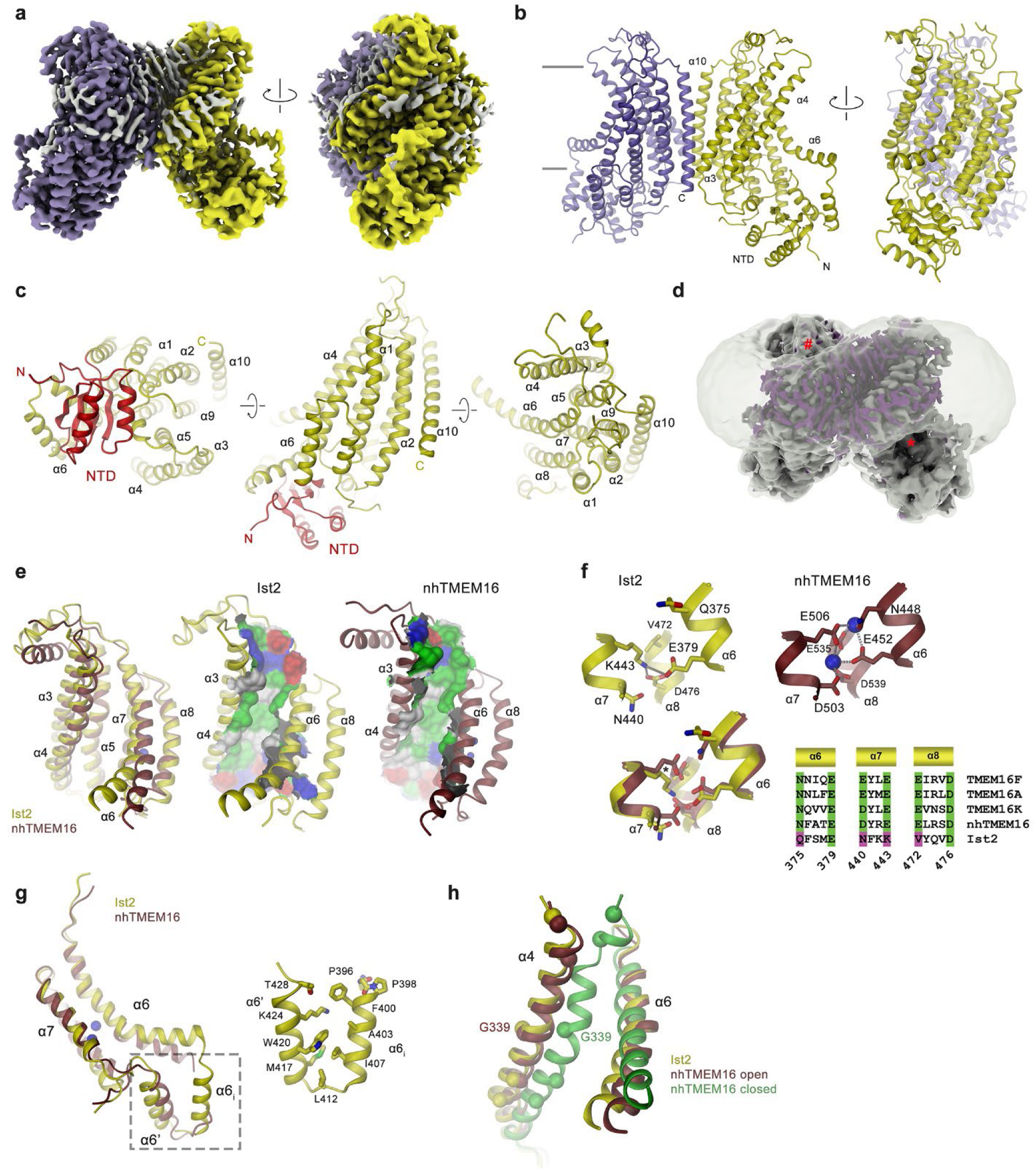
Ist2^716^ structure. **a**, Cryo-EM density of Ist2^716^ at 2.84 Å viewed from within the membrane. Relationship between views is indicated. Subunits are colored in yellow and violet, residual density presumably corresponding to detergent or lipids in grey. **b**, Ribbon representation of Ist2^716^ with colors and orientation as in **a**. Membrane boundaries are indicated, selected regions of the structure are labeled. **c**, Ribbon representation of the Ist2 subunit at indicated orientations with the NTD colored in red. Transmembrane helices are labeled. **d**, Cryo-EM density of Ist2^716^ with residual density corresponding to detergent and lipids colored in violet. A low-pass filtered map at low contour showing the distortion in the detergent micelle is superimposed. ‘#’ and ‘*’ indicate sites of deformation. **e**, Subunit cavity viewed from within the membrane along the long dimension of the dimeric molecule. Shown are α-helices 3-8 from a single subunit. Left, superposition of Ist2^716^ (yellow) and the Ca^2+^-bound open state of nhTMEM16 (brown, PDBID 6QM9), center, Ist2^716^, right nhTMEM16. Center, right, the molecular surface in the region of the subunit cavity is displayed and colored according to the physicochemical properties of contacting residues (polar, green, acidic red, basic, blue). **f**, Structure of the regulatory Ca^2+^ binding site in TMEM16 proteins and its equivalence in Ist2. Top, left, Ist2, right, nhTMEM16 with two Ca^2+^-ions bound. Bottom, left, superposition of equivalent regions of Ist2 and nhTMEM16, right, sequence alignment of relevant regions in fungal and murine TMEM16 proteins. The numbering refers to Ist2. **g**, Superposition of transmembrane α-helices 6 and 7 and their connecting region in Ist2 and nhTMEM16. Inset (right) shows blowup of the interacting cytoplasmic helices α6_i_ and α6’. E-G, Ca^2+^ ions bound to nhTMEM16 are displayed as blue spheres. **h**, Conformation of α4 and α6 lining the edge of the subunit cavity in a superposition of the subunits of Ist2, nhTMEM16 in presence (open, PDBID 6QM9) and absence of Ca^2+^ (closed, PDBID 8TPM). The protein is shown as Cα trace with the equivalent positions of α4 that are occupied by glycine residues in nhTMEM16 marked as spheres.

In the family members nhTMEM16 and afTMEM16, the absence of Ca^2+^ leads to a detachment of α6 from the binding site, which in turn triggers a conformational change in α4, leading to a collapse of the subunit cavity at its extracellular side^26,29^. In nhTMEM16, α4 contains several glycine residues distributed along the helix that affect its conformational properties^29^ (Fig. 2h). In Ist2, these glycines are not conserved, presumably increasing the stability of α4 in the observed conformation (Fig. 2h). Consequently, the cavity is found open in all determined structures of Ist2^716^, including the ones from nanodiscs (Extended Data Fig. 5e). However, even in case of Ist2, the stability of this open conformation is compromised by the interaction of a hydrophilic surface with the hydrophobic core of a detergent micelle. This is reflected in the structure of the membrane-inserted unit of full-length Ist2 in complex with Osh6 that was prepared in a different detergent mixture, where the cavity has rearranged to shield its hydrophilic surface by movements of α-helices 3, 4 and 6 (Extended Data Fig. 5h).

In summary, the structural and functional characterization of the membrane-inserted part of Ist2 has defined its function as a constitutively active lipid scramblase, exhibiting all hallmarks that are associated with a lowered energy barrier for lipid flip-flop.

### Properties of the C-terminal region of Ist2 and its interaction with Osh6

The absence of structured elements following the membrane inserted part of Ist2 (i.e. after residue 595) confirms previous predictions of the C-terminal linker as intrinsically disordered protein sequence^19^ (Figs. 1 and 2, Extended Data Figs. 1, 3, 4, Supplementary Fig. 1a). To obtain further insight into the properties of this region and its interaction with LTPs, we first determined the minimal sequence required for Osh6 binding by pulldown assays. To this end, we have initially based our construct design on previously published results, which suggested the core binding region to be located between residues 727 and 776 (ref. ^16,17^) (Supplementary Fig. 1a, c). This conserved stretch is situated close to the center of the 333 residues long flexible linker (excluding the very C-terminus tethered to the PM), measuring about 100 nm in a fully extended state and several times exceeding the 10-30 nm wide gap of the contact site between the ER and the plasma membrane. In our studies, we have systematically truncated the sequence on both ends of the binding site and found unperturbed interaction with a peptide encompassing residues 732 to 759, followed by a pronounced decline in shorter sequence stretches (Extended Data Fig. 6a). To determine the strength of the interaction, we quantified the binding of peptides comprising Ist2 residues 732-761 (Ist2^732–761^) and 732-756 (Ist2^732–756^) to Osh6 by isothermal titration calorimetry (ITC) and obtained in both cases affinities in the high nanomolar to low micromolar range (694.4 ± 54.2 nM and 714.0 ± 86.7 in case of Ist2^732–761^ and 1.1 ± 0.17μM in case of Ist2^732–756^, Fig. 3a, Extended Data Fig. 6b, c). To better understand the structural basis of the interaction, we have co-crystallized both peptides with an Osh6 variant lacking the first 30 amino acids (termed Osh6^ΔN^), which were disordered in previously determined crystal structures^10,11^. Data collected from complexes with either peptide extend beyond 1.9 Å resolution and provide a detailed view of the lipid transport protein in complex with its interacting region on the Ist2 C-terminus (Fig. 3b-g, Extended Data Fig. 6d-f, Extended Data Table 2). In both structures, Osh6 adopts a similar conformation as a curved beta sheet containing a single lipid binding site in its center that is closed by an N-terminal lid, as observed previously in complexes with the lipids PS and PI_4_P^10,11^. An exception concerns the conformation of the N-terminus, where α-helix 1 is extended, presumably as a consequence of crystal packing interactions (Fig. 3b, Extended Data Fig. 6d). The map obtained for the Osh6^ΔN^/Ist2^732–761^ complex shows well-defined density for residues 732-758 of Ist2, which bind to Osh6 on the opposite side of the lid covering the entrance to the lipid-binding pocket, burying 1’516 Å^2^ of the combined molecular surface (Fig. 3b-d and Extended Data Fig. 6e, f). The shorter peptide Ist2^732–756^ shows an equivalent binding mode and is defined until Arg 750 (Fig. 3c). The interacting residues on Osh6 are strongly conserved in homologs that are supposed to form similar complexes with Ist2-like proteins, but not in other family members that do not bind this tethering protein^16^ (Extended Data Fig. 6h, Supplementary Fig. 2). The N-terminal part of the peptide occupies a groove formed by residues 408-434 at the C-terminus of Osh6, that is stabilized by intermolecular backbone- and sidechain to backbone H-bonds. The latter include interactions of the sidechain of Tyr 391, Gln 417 and Thr 420 of Osh6 and Thr 736 of Ist2 (Fig. 3e, Extended Data Fig. 6f-h). Aliphatic contacts are pronounced between residues 737-742 containing two proline residues (Pro 738 and Pro 742), perturbing the extended conformation of the peptide in a region that is unusually hydrophobic compared to the remainder of the Ist2 linker (Fig. 3f, Extended Data Fig. 6f-h, Supplementary Fig. 1a, c). Further towards the C-terminus of the interaction region, the residues Ser 744-Arg 750 form a short helical segment, which is stabilized by an intramolecular hydrogen bond between the sidechain of Arg 750 and the backbone carbonyl of Thr 743, thereby inducing a conformation that is further stabilized by the stacking of the aromatic sidechains of Tyr 747 and Phe 751 (Fig. 3g, Extended Data Fig. 6f). Our results align with previously published data, in which mutation of Thr 736 and Thr 743 compromised the recruitment of Osh6 to contact sites^16^. Besides the observed stabilization of the peptide conformation, Arg 750 might also engage in long-range water-mediated interactions with the more distant Asp 141 of Osh6, a residue which has also been described to impair binding when mutated to alanine^16^ (Fig. 3g). Mutation of Arg 750 to Ala did indeed severely compromise binding to Osh6 in a pulldown assay and ITC, emphasizing the importance of this residue for the interaction (Extended Data Fig. 6i-k). Besides the detailed view of the interaction with its epitope on Ist2, the crystal structure also revealed well-resolved density of bound PS in the conserved binding pocket of Osh6, which was confirmed by mass spectrometry (Fig. 3h). Since Osh6 was overexpressed in *S. cerevisiae*, the lipid corresponds to the endogenous substrate and demonstrates that binding of PS and Ist2 are not mutually exclusive. Together, our results confirm the direct interaction between Osh6 and Ist2 through a conserved region in the center of the disordered tail involving residues 732-751 (Supplementary Fig. 1a, c).

**Figure 3.**
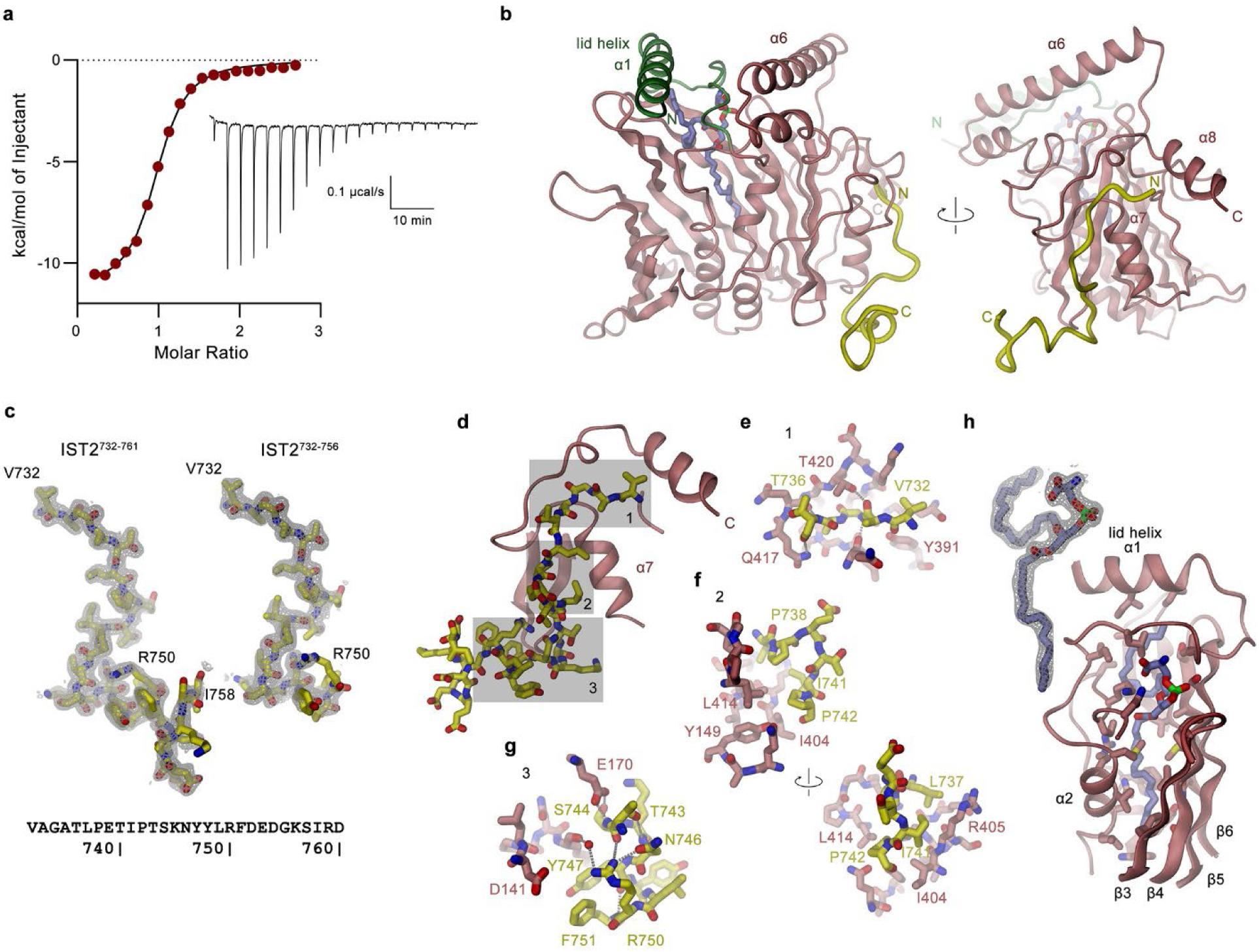
Interaction of Osh6 with the C-terminal region of Ist2. **a**, Binding of a peptide encompassing residues 732-761 of Ist2 (Ist2^732–761^) to Osh6. Binding isotherms were obtained from isothermal titrations of Ist2^732–761^ to Osh6. The data shows a fit to a model assuming a single binding site with the binding isotherm depicted as solid line with a K_d_ of 694.4 ± 54.2 nM. Errors represent fitting errors. The thermogram is shown as inset (right). **b**, Ribbon representation of the structure of the Osh6/ Ist2^732–761^ complex determined by X-ray crystallography at 1.9 Å. Ist2^732–761^ is colored in yellow, the N-terminal lid helix of Osh6 in green. A bound PS lipid is shown as sticks. **c**, 2Fo-Fc density of the refined structure superimposed on a model of the Ist2 binding region. Left, density from the Osh6/Ist2^732–761^ complex defining the conformation of residues 732-758 of the peptide, right, density of the Osh6 complex with the shorter Ist2^732–756^ peptide determined at 1.9 Å defining the conformation of residues 732-750 of the peptide. The sequence of the Ist2^732–761^ peptide is shown below. **d**, Sections of the Osh6-Ist2 interactions revealed in the Osh6/Ist2^732–761^ complex. A model of the defined parts of the Ist2^732–756^ peptide in stick representation with its interacting region of Osh6 represented as ribbon. Different regions of the interaction are indicated as shaded area and shown as blow-up with Osh6 represented as sticks in panels e-g. (**e**-**g**) Sections of the Osh6-Ist2 interactions highlighted in d in relation to the Osh6 binding sequence on Ist2. **e**, N-terminal region, **f**, center and **g**, C-terminal region. **h**, 2Fo-Fc density of the bound PS lipid (left) and binding lipid binding region of Osh6 with interacting sidechains shown as sticks.

### Osh6 interaction with membranes

After characterizing its binding to Ist2, we have investigated the interaction of Osh6 with lipid bilayers, following up on earlier studies, which described this process as critical step during the extraction and release of its lipid cargo^31^. To monitor its avidity for negatively charged lipid patches, we mixed fluorescently labeled Osh6 with lipid nanodiscs composed of the anionic lipid POPG and the neutral lipid POPE and were able to detect its co-elution with the nanodisc fraction in size-exclusion chromatography (Fig. 4a). As the binding of its specific substrates PS or PI_4_P decreases its affinity to lipid bilayers, we subsequently overexpressed and purified Osh6 from *E. coli*, whose membranes have a low PS content, and confirmed the absence of bound lipids by mass spectrometry. With this sample, we proceeded with a structural characterization by cryo-EM (Fig. 4b, Extended Data Fig. 7). In 2D reconstructions of this dataset, distinct classes show the density of single Osh6 molecules bound to the edge of a lipid nanodisc (Extended Data Fig. 7b). After intensive classification, a 3D reconstruction permitted the placement of Osh6 with its loop region leading to the PS binding site in presumed contact with lipids (Fig. 4c, d). In this binding mode, the Ist2 binding epitope is located on the opposite face of the protein remote from the membrane, permitting simultaneous binding to its tether (Fig. 4d). The location of Osh6 at the edge of nanodiscs illustrates its preference for distorted membrane regions as no density was found in the center of the disc, neither did we detect interactions with the isolated scaffold protein. This preference is remarkable considering the observed membrane distortion by the transmembrane part of Ist2 (Fig. 2d, Extended Data Fig. 5d, e).

**Figure 4.**
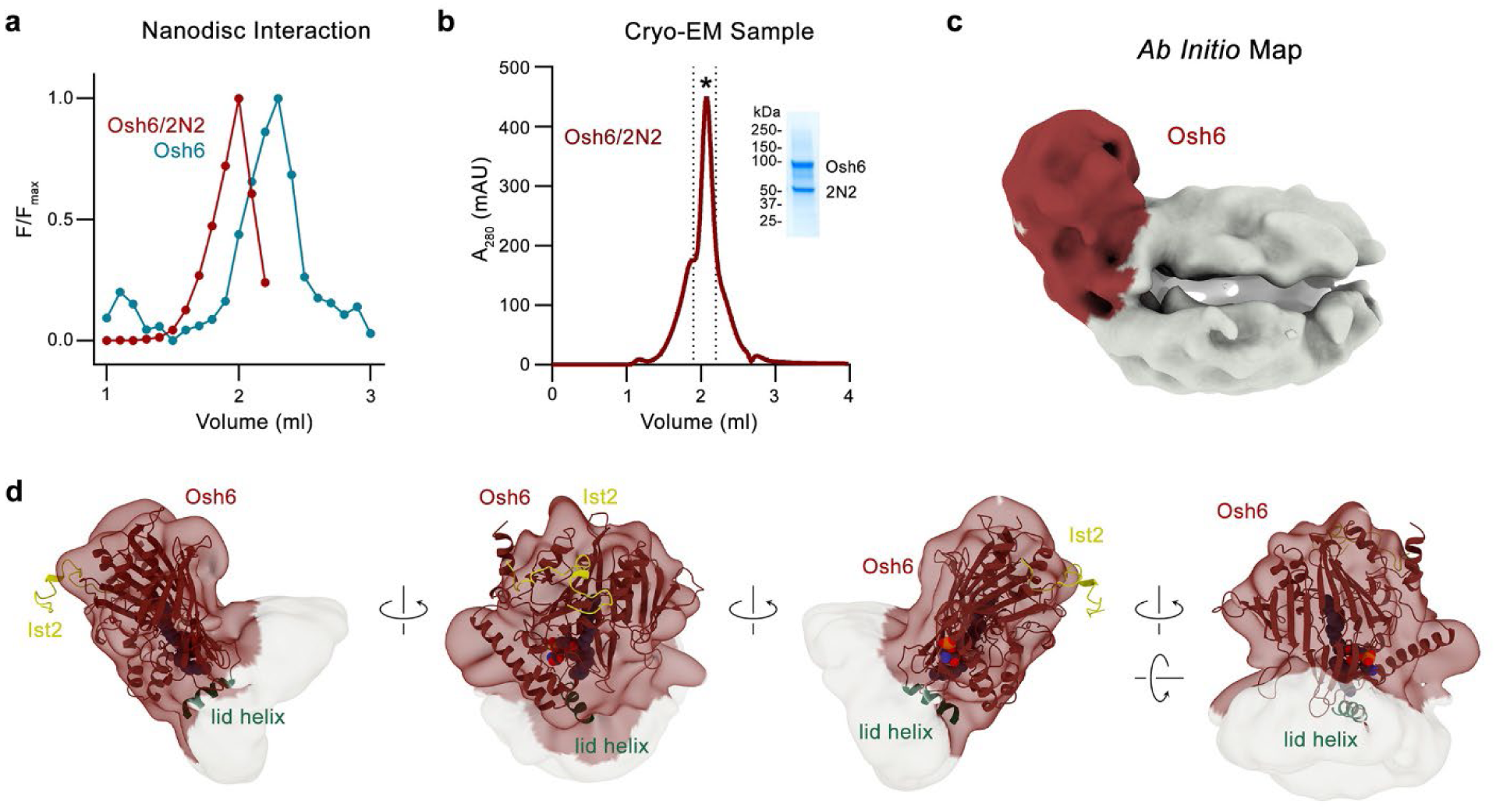
Structural properties of the Osh6 interaction with lipid membranes. **a**, Size exclusion chromatograms of Osh6 in absence and presence of 2N2 lipid nanodiscs monitored by the fluorescence of the labeled LTP. The co-migration with nanodiscs is reflected in the decreased elution volume. **b**, SEC profile and SDS PAGE gel (inset) of the Osh6/2N2 lipid nanodisc sample used for cryo-EM analysis. **c**, Ab initio reconstruction of Osh6 (red) bound to the edge of a lipid nanodisc (white) obtained from a cryo-EM dataset. **d**, Model of Osh6 fitted into the masked and refined cryo-EM density. The lid helix α1 is colored in green. The interacting Ist2 peptide (yellow), which was not included in the sample, is displayed for reference. The bound POPS is shown as space filling model, the relationship between views is indicated.

### Cellular assays suggest continual binding of Osh6 to Ist2 during lipid transfer

After characterizing the binding of Osh6 to Ist2 on a biochemical and structural level, we have investigated the role of the Ist2 linker and its interaction with Osh6 for lipid transport. In our studies, we were particularly interested in the influence of linker length, the impact of the relative location of the binding site and the question of whether Osh6 remains bound to the cytoplasmic tail of Ist2 during lipid transfer. The comparably long interaction sequence and the consequently high binding affinity suggest a stable interaction of both proteins (Fig. 3a-d), while the length of the linker, several times exceeding the distance between adjacent membranes in the contact site, and the location of the binding site close to its center would render the shuttling of the bound protein between membranes of opposite compartments a possible scenario.

To investigate this hypothesis, we employed a previously established cellular assay, that is based on the rescue of a growth defect in an *ist2/psd1* KO strain, as a consequence of PC limitation^17^. The described growth deficit can be alleviated either by transformation with a construct coding for wild-type Ist2 or by supplementation of choline, a PS precursor of the Kennedy-Pathway (Fig. 5a, Extended Data Fig. 8a-c). In contrast, growth is not rescued by overexpression of the truncated construct Ist2^716^, which does not contain the Osh6 binding site, or the longer construct Ist2^921^ including the binding site but lacking the C-terminal cortical localization sequence, which both were previously shown to mislocalize to the perinuclear ER^32^ (Fig. 5a, Extended Data Fig. 8c, Supplementary Fig. 3a). Using this assay, we assessed the ability of modified Ist2 constructs to form a functional lipid transport system at MCSs in yeast. We first probed the effect of an alteration of the Osh6 binding site by either introducing the point mutation R750A, which strongly weakened binding (Extended Data Fig. 6i-k), or by scrambling the sequence of the binding site without altering its composition (Supplementary Fig. 3b). In both cases, we find a severely compromised complementation of growth, which is already pronounced in the point mutation and completely abolished in case of the scrambled binding site, underlining the role of the interaction for lipid transport (Fig. 5b, Extended Data Fig. 8d). We then probed the impact of the relative location of the binding site on the linker by moving the site by 83 residues towards either the N- or C-terminus of the protein while preserving its overall length (Supplementary Fig. 3c). In this case, we found a severely reduced growth phenotype, which is more pronounced upon movement of the site towards the N-terminus than towards the C-terminus (Fig. 5c and Extended Data Fig. 8c) as expected from the shorter length of the respective linker region.

**Figure 5.**
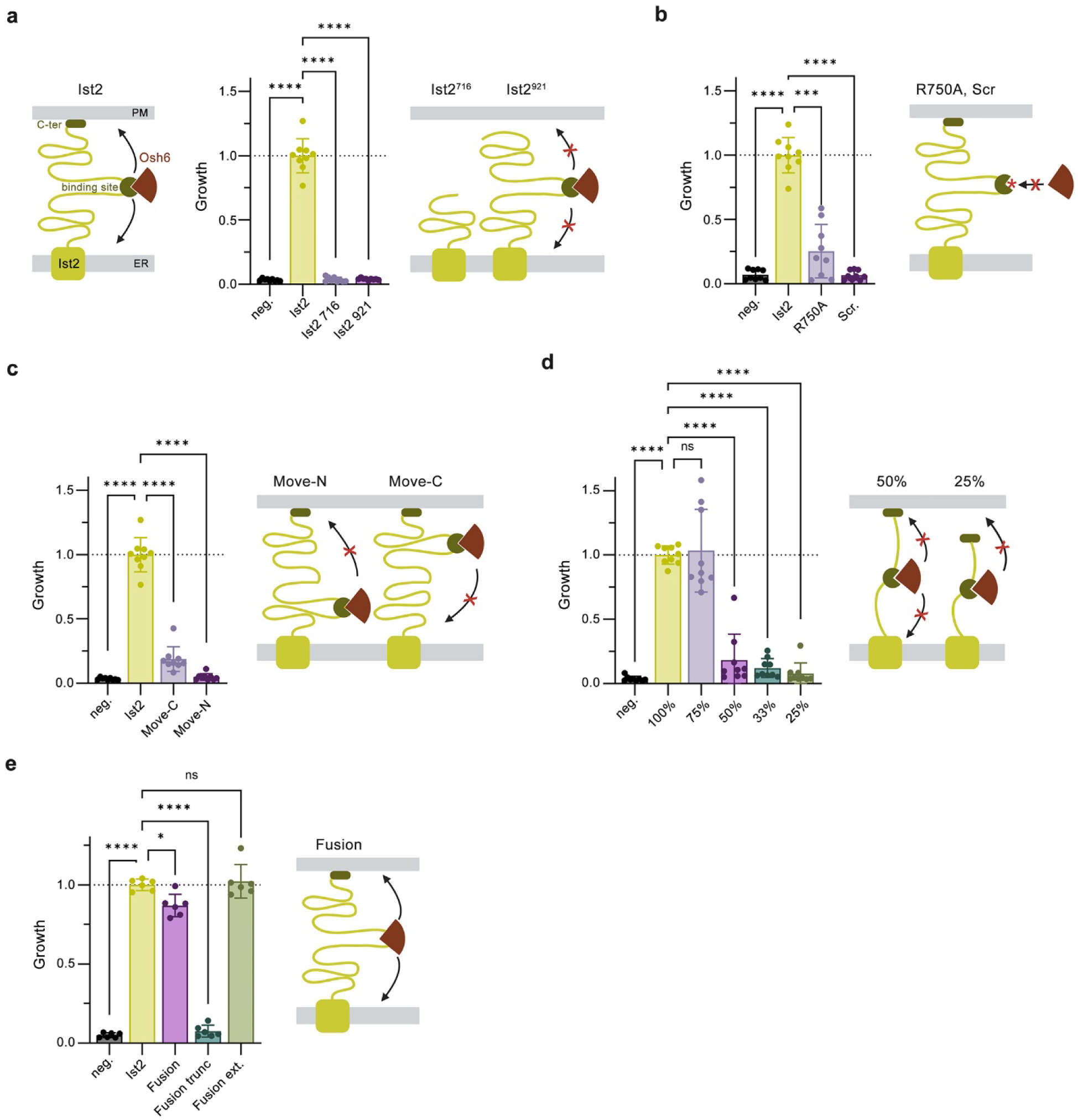
Functional properties of Ist2 constructs probed in yeast complementation assays. Cell growth assayed upon overexpression of Ist2 constructs in a *S.cerevisiae ist2/psd1* KO strain. **a**, Comparison of WT to the constructs Ist2^921^ lacking 25 residues of the C-terminus including the PM binding region and Ist2^716^ lacking 230 residues including the Osh6 binding site. **b**, Comparison of WT to a protein carrying the mutation R750A in the Osh6 binding site of Ist2, which strongly reduces the interaction, and another construct where the sequence of the binding site was scrambled without changing its composition. **c**, Comparison of WT to constructs where the Osh6 binding site was moved by 83 residues towards the N- and C terminus of the protein. **d**, Comparison of WT to constructs where the Ist2 linker was symmetrically shortened by removal of residues from the center of both sequence regions N- and C-terminal to the binding site to reach the indicated fraction of its original length. **e**, Comparison of WT to constructs with the Osh6 binding site removed and Osh6 fused into the equivalent position into the linker (Fusion), a fusion construct where the linker was symmetrically truncated by 50% in the center of both regions flanking the fused LTP (Fusion trunc) or expanded by 50% by introducing a Gly-Ser peptide sequence (Fusion ext.). **a**-**e**, Schemes illustrate the properties of Ist2 constructs. Bars display mean normalized growth compared to WT, n=3x3 replicates (2x3 replicates for Fusion tunc and Fusion ext), errors are s.e.m., values of individual experiments are shown as dots. The statistical significance is indicated.

Next, we have investigated the effect of linker length on complementation upon shortening of the C-terminus by a symmetric removal of residues from the center of both regions flanking each side of the Osh6 binding sequence (Supplementary Fig. 3d). Assuming complete flexibility of the linker on both sides of the binding site, its length would exceed the experimentally observed distance between the plasma membrane (PM) and the ER at least by a factor of two^19^, which would permit a shuttle mechanism of Osh6 between both membranes while remaining bound to Ist2. In our studies, we observe pronounced complementation in constructs containing up to three quarters of the residues and a sharp decline in shorter constructs with an apparent minimal length of between 170-250 residues suggesting that shorter constructs would be insufficient to permit bound Osh6 to reach both membranes (Fig. 5d, Extended Data Fig. 8e). Finally, we have investigated a fusion construct where we have removed the Osh6 binding site and instead covalently inserted the protein at the same position into the linker (Supplementary Fig. 3e). Remarkably, this construct, which is biochemically well behaved, allowed nearly WT-like complementation of the growth phenotype, which we found unaltered upon extension and considerably reduced upon shortening of the linker (Fig. 5e, Extended Data Fig. 7f, g).

Taken together, the results from the yeast complementation assays demonstrate the importance of the Osh6/Ist2 interaction and its positioning in the center of a long flexible linker to permit shuttling of the lipid binding protein between opposed membranes of the contact site and thus strongly suggest that the complex remains intact during lipid transfer.

## Discussion

As the main hub for lipid synthesis in eukaryotes, the ER plays an important role in the distribution of lipids to other cellular compartments. In recent years, non-vesicular lipid transport at MCS of the ER has gained considerable attention^33^ and was shown to proceed via two general mechanisms. Channel-like transport processes, which are mediated by rigid conduits that bridge the membranes of contacted compartments, were described for the Golgi-localized protein VPS13 (ref. ^34,35^), the autophagy-related protein ATG2 (ref. ^36,37^) and the ERMES complex mediating contacts between the ER and mitochondria^38,39^ (Fig. 6a). These proteins harbor a continuous hydrophobic permeation path facilitating the diffusion of lipids across the contact site with low specificity.

**Figure 6.**
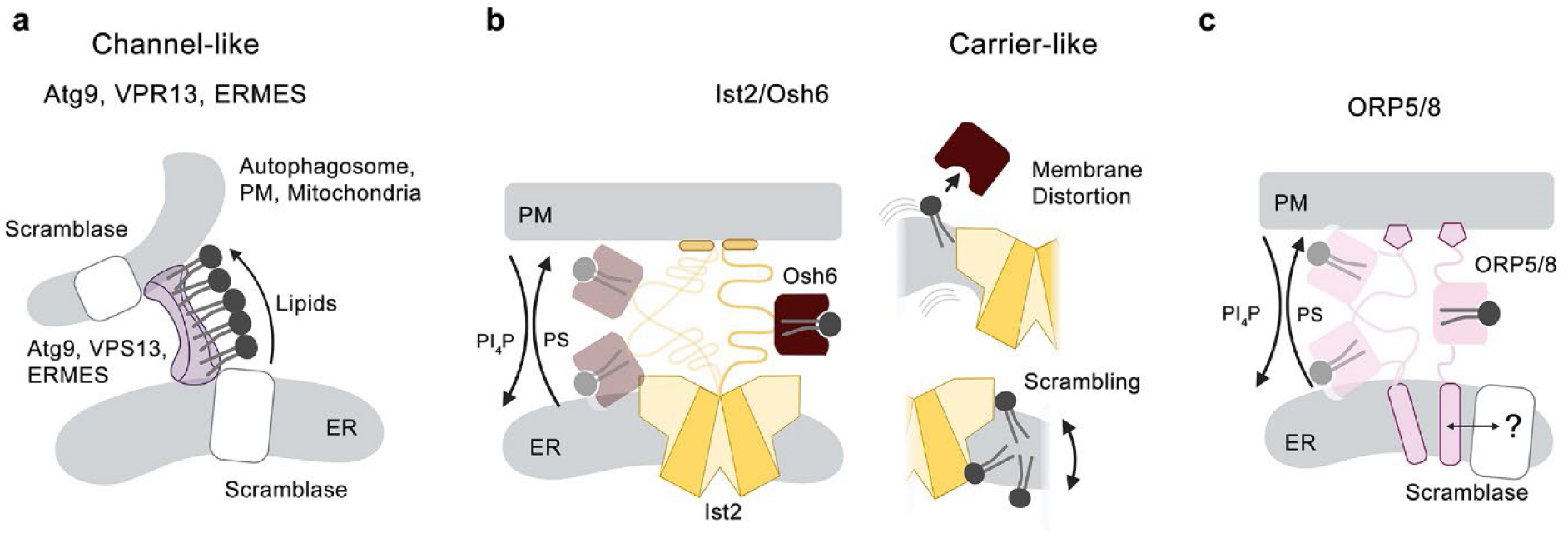
Mechanisms of non-vesicular lipid transport at membrane contact sites. **a**, Channel-like mechanism as observed in the proteins Atg9 and VPR13 provide a continuous hydrophobic environment for the diffusion of lipids between connected leaflets of contacted membranes. Interacting lipid scramblases equilibrate the imbalance between leaflets upon extraction and insertion of lipids. **b**, **c**, Carrier-like mechanisms use lipid transfer proteins (LTPs) that are tethered to a flexible linker bridging both membranes of the contact site as in case of the Ist2/Osh6 system (**b**) or the mammalian homologs ORP5 and 8 (**c**). The LTPs carry a binding site that is specific for the lipids PS and PI_4_P. In case of Ist2/Osh6, the lipid scramblase function of Ist2 might distort the membrane and thus facilitate lipid extraction and insertion and equilibrate imbalances due to the insertion of lipids with large headgroups. In case of ORP5 and 8 the interaction with a lipid scramblase is still hypothetical. **a**-**c** Figures were created using BioRender.

A second mechanism mediates selective transport via carrier proteins, such as members of the ORP family, which contain a specific and saturable lipid binding site^7,40^. These proteins are tethered to a flexible linker that is attached to a membrane-embedded domain, which localizes the lipid carriers to the MCS to allow the shuttling between both compartments to take up and release their cargo at the respective target membranes^41,42^ (Fig. 6b, c). The tether can either be covalently attached to the lipid carrier, as in case of certain mammalian ORP homologs, or be a distinct protein as in case of the yeast Ist2/Osh6/7 system^40,43^.

In our study, we have investigated the Ist2/Osh6 system in *S. cerevisiae* with a focus on the role of the membrane protein Ist2, a homolog of the TMEM16 family that encompasses ion channels and lipid scramblases^22^. We were particularly interested in the structural properties of Ist2 and whether it has retained the lipid transport activity found in other family members. Although its ability to scramble lipids was challenged previously^24^, we have here confirmed this function by *in vitro* lipid transport assays and a thorough structural characterization, which together provide a mechanistic explanation for its constitutive activity (Figs. 1e-h and 2). In our data, Ist2 shows the structural hallmarks of a TMEM16 lipid scramblase with unique features in the catalytic subunit cavity region that lead to a stabilization of the ‘open’ conformation (Fig. 2e-h). Similar to structures obtained for the Ca^2+^-bound active form of its fungal homologs afTMEM16, nhTMEM16 and the mammalian TMEM16K^23,26,27^, this cavity exposes a hydrophilic surface to the membrane (Fig. 2e). Unlike in most family members, the regulatory Ca^2+^ binding site is altered in Ist2 and its close homologs in *Saccharomycetaceae* to stabilize the observed conformation even in the absence of Ca^2+^ (Fig. 2f, Supplementary Fig. 1b). Since this binding site is highly conserved in the TMEM16 family, its absence raises the question of whether Ca^2+^ regulation might have emerged from an unregulated ancestor and subsequently evolved from weakly regulated scramblases in intracellular organelles to the tightly regulated family members found in the plasma membrane of higher eukaryotes, where the suppression of basal activity is critical to prevent severe physiological consequences^28,44^.

The identification of Ist2 as a passive lipid transporter aligns with the proposed role of lipid scramblases as essential components of non-vesicular lipid transport systems^45^. Their relevance is evident in the case of channel-like systems, where a net lipid flux between two compartments creates tension through lipid extraction from a single leaflet, which can be relaxed by a scramblase. In case of carrier-like processes, the benefit of a lipid scramblase is less obvious, since transport proceeds as exchange of counter-transported lipids. PS transport from the ER to the plasma membrane is by Osh6 compensated by PI_4_P transport in the opposite direction preventing a pronounced change in the membrane surface of either compartment^11^. However, even in this case scrambling might be necessary to relax local lipid imbalances arising from the accumulation of the bulky inositol headgroup of PI_4_P in the vicinity of Ist2. Additionally, our data hints at another scenario, in which the distortion of the local membrane environment by Ist2 may facilitate the access of lipids to the Osh6 binding site. In both bulk and carrier models of lipid transfer, lipid extraction from the bilayer has been described as the rate limiting step of the process^34,46^. Local perturbation of the membrane could consequently help to lower this energy barrier. In our cryo-EM studies of Osh6-nanodisc complexes, we observe binding of Osh6 to the periphery of the lipid bilayer (Fig. 4d, e), indicating a preference of Osh6 for curved membrane regions. The role of the protein-induced membrane deformation to facilitate lipid extraction by Osh6 could be particularly relevant in light of the cellular localization of Ist2 at flat ER-sheets observed in cryo-CLEM studies, where the global membrane curvature is minimal^13^.

Besides the characterization of its transmembrane domain, we have investigated the unstructured cytosolic region of Ist2, which functions as a tether at ER-PM contact sites that recruits the soluble Osh6 via a conserved binding site that is located in its center. As noted previously, the linker region is of sufficient length to span the experimentally observed distance between the two membranes several times^16,19^. We therefore explored the hypothesis that Osh6 remains bound to Ist2 during lipid transport. Our results reveal the structural basis of the interaction with the Ist2 C-terminus, showing that its affinity is strong enough to support sustained attachment to the linker in a cellular environment (Fig. 3a-g and Extended Data Fig. 6). Cellular complementation assays further demonstrate that this persistent binding of Osh6 is crucial for efficient lipid transport (Fig. 5), despite the intrinsic membrane avidity of Osh6 observed *in vitro* (Fig. 4). The permanent attachment of Osh6 to Ist2 during lipid transport is reminiscent of the homologous OSBP-related lipid transport systems found in mammals^47^, where the PS/PI_4_P exchangers ORP5 and ORP8 directly act as membrane tethers through a N-terminal PH domain, which binds to PIPs in the PM, and a C-terminal transmembrane helix, which anchors the protein to the ER^43,48^. Both systems share equivalent features of a lipid carrier that is anchored to a flexible tether like a bead on a string (Fig. 6b, c). From a biophysical perspective, immobilization of the soluble transport protein is beneficial as it decreases its accessible volume and essentially limits its movement to one-dimensional diffusion. The tethering also preserves the localization of the lipid binding protein at the contact site, avoiding its diffusion into the cytoplasm. Together, these factors can enhance the rate of lipid exchange and compensate for the comparably low transport capacity of a single lipid molecule per carrier.

Since the covalent association of a membrane tether to a lipid scramblase is a unique feature of *Saccharomycetaceae*, while ORP systems in higher eukaryotes only contain a simple membrane anchor, it will be interesting to investigate whether these proteins recruit accessory scramblases to take over the role of Ist2 in mammalian systems. Interestingly, the human homologues TMEM16K and TMEM16H have been shown to localize to MCSs between the ER and endosomes, although their exact function has remained elusive^49,50^. While previously published results suggest that the TM unit of Ist2 is to a large part dispensable^17^, our data clearly demonstrate that the ER domain exerts a functional role beyond anchoring.

In summary, our results have provided detailed mechanistic insight into lipid transport at MCS. Whereas channel-like lipid transport systems have previously been investigated in structural and mechanistic detail, tethered carrier systems have mainly been studied from a cell-biological perspective. Unlike the former, which transport lipids in bulk and with low selectivity, lipid carriers are unique in their strong lipid specificity, which has been demonstrated previously in *in-vitro* systems^11^. Our data now extends the previous view of Ist2 as a simple membrane tether as we show that it acts as a lipid scramblase and that the continued binding of Osh6 is indispensable for its efficiency as a lipid transporter. While more detailed experiments will be necessary to better understand the interplay between both parts of the protein, our study opens up a compelling view on the role of intracellular TMEM16 proteins at MCS and it demonstrates the association of an active scramblase with a carrier-like lipid transport system.

## Acknowledgements

This research was supported by grants from the Swiss National Science Foundation (No. 310030_215236 to R.D.). Cryo-EM data were collected at the Center for Microscopy and Image Analysis (ZMB) of the University of Zurich. We thank Piotr Szwedziak for support during microscope operation, Beat Blattman and Görkem Kurduldu of the Protein Crystallization Center at UZH, for their support with crystallization. X-ray crystallography data was collected at the Beamline P13 at DESY (Hamburg). The support of Francesco Nai and Rajiv Bedi and Staff of the P13 beamline during Data collection is acknowledged. We thank Benedikt Wimmer for advice regarding Cryo-EM data processing. The yeast strains used for complementation experiments were generously provided by Prof. Christopher J. R. Loewen (University of British Columbia). Members of the Dutzler group are acknowledged for their help at various stages of the project.

## Author Contributions

M.A. expressed and purified proteins, collected and analyzed cryo-EM and X-ray data, carried out lipid transport and yeast complementation assays. A. S. has established the yeast complementation assay performed ITC measurements and supported M.A. in various experiments. M.A. and R.D. jointly planned experiments, analyzed the data and wrote the manuscript.

## Methods

### Cloning of expression constructs

All expression constructs were generated by FX-cloning^51^ unless specified otherwise. For yeast expression constructs, the sequence of nhTMEM16, Ist2 full length (Ist2fl) and Ist2 containing residues 1-716 (Ist^716^) was cloned into a modified FX-compatible pYES2 vector with an N-terminal SBP-myc fusion followed by a 3C recognition site (pYESnSM3). Ist2 tail variants used in the yeast complementation assay (Supplementary Figure X) were either ordered as synthetic DNA constructs from Genscript (Ist2^50%^, Move-C, Move-N, Ist2-Osh6 Fusion) or cloned via a seamless exchange of synthetically ordered tail fragments into a ccdb cassette flanked by SapI cleavage sites introduced between residues 590-591 (serine) and 926-927 (alanine) in the full-length construct (Ist2^25%^, Ist2^33%^, Ist2^75%^, Ist2-Osh6 Fusion extended, Ist2-Osh6 Fusion truncated). The sequence of Osh6 and Osh6Δ1-30 was cloned into pYESnSM3. Point mutations Ist2_R750A and Osh6_C389S were introduced using a modified QuickChange protocol^52^. For bacterial expression constructs, the sequence of Osh6 was cloned into a pGEX-4T vector using BamHI and EcoRI. The forward primer was extended in the 5’ end introduce an N-terminal 3C cleavage sequence following the Thrombin recognition site. TMEM16A was inserted into a pcDXc3GMS vector using FX cloning^53^. The sequence for MSP 2N2 and MSP E3D1 were purchased from Addgene. All constructs were verified by sanger sequencing.

### Protein expression and purification

Recombinant Ist2 and nhTMEM16 constructs were transformed into the *S. cerevisiae* expression strain FGY217 using the LiAc/SS carrier DNA/PEG method^54^. Positive clones were grown in SC-Ura selection medium at 30 °C in a fermenter until and OD_600_ of 0.7 was reached. Protein expression was induced by addition of 2% galactose and the temperature was reduced to 25 °C. After 40 h, the cells were harvested at 4000 g for 10 min. For Ist2 1-716 and nhTMEM16, cells were resuspended in Buffer A (20 mM HEPES pH 7.6, 150 mM NaCl) and supplemented with 1mM MgCl_2_, protease inhibitors (PMSF, Benzamidine, Pepstatin A, Leupeptin), DNAse and Lysozyme and lysed in a HPL6 high pressure cell disruptor at 40 kpsi. Cell debris was removed by centrifugation at 6,000 g for 30 min at 4 °C Membranes were harvested from the supernatant by ultracentrifugation at 150,000 g for 1 h at 4 °C. The membrane pellet was resuspended in minimal volumes of buffer A using a dounce-homogenizer, flash frozen in liquid nitrogen and stored at -80 °C until further use. For protein purification, membranes were resuspended in 5x pellet volume of Buffer A supplemented with protease inhibitors, 5% glycerol and 2% GDN (glyco-disogenin, Anatrace). Extraction was carried out for 1.5 h at 4 °C. The insoluble fraction was removed by ultracentrifugation at 85,000 g for 1 h and the solubilized protein in the supernatant was bound in batch to Streptactin resin (IBA) for 2 h at 4 °C. Resin was washed with 60 column volumes (CV) of Buffer B (20 mM HEPES pH 7.6, 150 mM NaCl, 5% Glycerol, 0.03% GDN) using gravity flow and bound protein was eluted in 5 CV of Buffer B supplemented with 5 mM d-Desthiobiotin. Protein was concentrated to 2 mg ml^-1^ and loaded onto a Superose 6 10/300 GL column pre-equilibrated with Buffer C (20mM HEPES pH 7.6, 150mM NaCl, 0.03% GDN). Peak fractions were pooled and concentrated to 1-2.5 mg ml^-1^. Purification of Ist2fl was performed similarly with the exception of Buffer A consisting of 40 mM HEPES pH 7.6, 500 mM NaCl, 5% Glycerol and GDN being replaced by 2% Tridecylmaltoside:CHS at a molar ratio of 9:1 during extraction. Purification of TMEM16A for reconstitution was performed similarly to Ist2 1-716, but in the presence of 2mM EGTA to avoid Ca^2+^-binding and the GFP tag was cleaved using 3C protease. Murine TMEM16A was expressed in HEK293 GnTI^-^ cells grown in Hyclone media with 1% Penicillin Streptomycin, 1% FBS, 2% Glutamine and 0.15% Kollophor p188 via transient transfection in suspension culture. The expression vector was mixed with PEI MAX in a 1:2.5 ratio and valproic acid was added to a final concentration of 3.5 mM after transfection. The protein was expressed at 37 °C for 48 h. Cells were harvested via centrifugation and washed once in PBS. TMEM16A was extracted in 20 mM HEPES pH 7.4, 150 mM NaCl, 2 mM EGTA and protease inhibitors (see above) by addition of 2% GDN for 2 h at 4 °C and the insoluble fraction was removed by ultracentrifugation at 85,000 g for 30 min at 4 °C. The supernatant was bound in batch to Streptactin resin (IBA) for 2 h at 4 °C and washed with 60 CV of SEC buffer (20 mM HEPES pH 7.4, 150 mM NaCl, 2 mM EGTA, 0.03% GDN). Bound protein was eluted in wash buffer containing 5 mM d-Desthiobiotin and concentrated to 1.5 mg/ml. The c-terminal SBP-myc-GFP tag was cleaved by addition of 0.1 mg/ml 3C protease for 45 min at 4 °C. The cleaved protein was loaded onto a Superose 6 10/300 GL column equilibrated in SEC Buffer and peak fractions were pooled and concentrated to 1 mg/ml. Recombinant Osh6 was purified from *S.cerevisiae* (for crystallization, Osh6Δ1-30, and ITC experiments, WT Osh6) and from *E.Coli* (for cryo-EM experiments and pulldowns). In *S.cerevisiae*, Osh6 was expressed similar to Ist2. Cells were resuspended in Buffer D (40 mM HEPES pH 7.6, 150 mM NaCl, 1mM MgCl_2_ and 0.5 mM TCEP) supplemented with protease inhibitors, DNAse and lysozyme and lysed with a HPL6 at 40 kpsi. The cytosolic fraction was cleared by ultracentrifugation at 150’000 g for 1 h and bound to Streptactin Resin (IBA) for 2 h as described above. The resin was washed with 60 CV of buffer E (25 mM HEPES pH 7.6, 150 mM NaCl, 0.5 mM TCEP) and eluted with Buffer E supplemented with 5 mM d-Desthiobiotin. The protein was concentrated to 3 mg/ml and loaded onto a Superdex200 10/300 GL column equilibrated in buffer E. Peak fractions were pooled, supplemented with 5% glycerol, concentrated, aliquoted and flash-frozen. Recombinant Osh6Δ1-30 was purified similarly, except that the pH of all buffers was adjusted to 7.8 and TCEP was omitted from the final gel-filtration step. For Osh6 expressed and purified in *E. coli*, BL21 transformants were grown in LB at 37 °C. Upon reaching an OD_600_ of 0.8, the temperature was reduced to 25 °C and gene expression was induced by addition of 0.3 mM IPTG at an OD_600_ of 1.2. The protein was expressed over night at 18 °C. Cells were harvested, resuspended in Buffer D and lysed by sonication. The insoluble fraction was removed by ultracentrifugation at 150’000 g, for 1 h at 4 °C and the expressed protein in the supernatant was bound in batch to GSH-Agarose (Genscript) for 1.5 h at 4 °C. The resin was washed with 50 CV of buffer E and 50 µM 3C protease was added to cleave protein from the resin for 45 min at 4 °C. The flow-through containing the protein was concentrated and loaded onto a Superdex200 10/300 GL column equilibrated in buffer E.

### Lipid Preparation and Preparation of Lipid Nanodiscs

All lipids were purchased from Avanti and mixed in Chloroform at the desired molar ratio, dried under nitrogen and washed in diethylether. After forming a thin lipid film under nitrogen, residual traces of solvent were removed in a desiccator overnight. The film was rehydrated in SEC Buffer (25 mM HEPES pH 7.4, 150 mM NaCl) for 2 h at room temperature (RT), sonicated briefly, subjected to three freeze-thaw cycles, aliquoted and stored at -80 °C. For Nanodisc reconstitution of Ist2^716^, the purified protein was concentrated to 0.87 mg/ml and reconstituted using the scaffold protein MSPE3D1 and a POPC:POPG 3:1 lipid mix at a protein:lipid ration of 1:50 with a 5x excess of empty nanodiscs. Protein and lipid mix were incubated in Buffer C for 30 min at 4 °C. Scaffold protein was added and incubated for another 30 min. The mixture was added to clean SM-II Biobeads at a ratio of 20 mg Biobeads/mg of detergent. The detergent was removed overnight at 4 °C by gentle agitation. Excess empty nanodiscs were removed by size exclusion chromatography on a Superose 6 10/300 GL column. The fractions containing the reconstituted protein were pooled and concentrated to 1.1 mg ml^-1^.

Empty nanodiscs for Osh6 binding were formed in SEC Buffer using a 1:200 protein:lipid ratio of MSP2N2 and a 3:1 POPC:POPG lipid mix. Nanodiscs were reconstituted as described above. The detergent-free supernatant was concentrated to 3.6 mg ml^-1^ and incubated with a 1.2x excess of purified Osh6 at a final concentration of 28 µM. The sample was injected multiple times on a Superose 6 5/150GL column equilibrated in SEC Buffer. The peak fractions were pooled and concentrated to 1.63 mg ml^-1^. Purity of the sample was assessed by SDS PAGE.

### Protein reconstitution and lipid scrambling assay

For lipid scrambling experiments, Ist2^716^, nhTMEM16 and TMEM16A were reconstituted into proteoliposomes using liposome destabilization as described previously^55^. In brief, liposomes at 20 mg ml^-1^ in Resuspension Buffer (20 mM HEPES pH 7.5, 300 mM KCl, 2 mM EGTA) composed of a 8:2 mixture of Soybean Polar Extract:Cholesterol with 0.5 mol% tail-labeled NBD-PE (Avanti) were destabilized by adding 0.02% aliquots of Triton-X-100. The onset of liposome destabilization was observed by the decrease of scattering at 540 nm in a UV-Vis-spectrofluorometer. Protein was added at a protein:lipid ratio of 1:300 (nhTMEM16) or 1:200 (Ist2^716^, TMEM16A) to account for different reconstitution efficiencies. For empty liposomes, Buffer C was added at an equivalent volume. Detergent was removed by four sequential additions of 20 mg SM-II Biobeads/mg of lipids over night at 4 °C and liposomes were harvested at 170,000 g for 30 min. The liposome pellet was resuspended to a final lipid concentration of 10 mg/ml, subjected to 3 freeze-thaw cycles and stored at -80 °C until further use. For the assay, liposomes were diluted to a final concentration of 20 µM in Assay Buffer (80 mM HEPES pH 7.5, 300 mM KCl, 2 mM EGTA) in a quartz glass cuvette. After recording the fluorescence of NBD for 60 s, 30 mM Dithionite was added to bleach the fluorescence of NBD in the outer leaflet of the liposomes. The fluorescence was recorded for an additional 6 minutes. Each fluorescence trace was normalized to the NBD fluorescence level right before the addition of dithionite. Statistical analysis was performed using ANOVA in GraphPad Prism. Because the final plateau differs depending on the reconstitution efficiency, for a statistical analysis samples reconstituted from the same batch of destabilized liposomes on the same day were matched in the analysis.

### Pulldown assays

Constructs coding for N-terminally GST tagged peptides of different length encompassing the Osh6 binding epitope of Ist2 were transformed into *E. coli* BL21 cells. Bacteria were grown in LB medium until an OD_600_ of 0.8 and gene expression was induced upon addition of IPTG to a final concentration of 0.3mM. Expression was carried out at 25 °C for 2.5 h. Cells were harvested, washed in PBS and resuspended in Lysis Buffer (25 mM HEPES pH 7.5, 300 mM NaCl, 10% Glycerol w/v, 0.5 mM TCEP, 0.1% Triton, 1 mM MgCl_2_) supplemented with Protease Inhibitors and DNAse. Cells were lysed by sonication and the insoluble fraction was removed by centrifugation at 150.000 g for 1 h at 4 °C. The cleared lysate was frozen in liquid nitrogen and stored at -80 °C until further use. The thawed lysate was diluted 1:10 in binding buffer (25 mM HEPES pH 7.4, 150 mM NaCl, 0.5 mM TCEP, 1 mg/ml BSA), mixed with Glutathione agarose (Genscript) at a final ratio of 1:4 and incubated for 2 h at 4°C under gentle agitation. Beads were washed 5x with a 10-fold excess of binding buffer to remove unbound proteins. 20 µl of coated beads were incubated with a final concentration of 5 µM Osh6 in a total volume of 40 µl for 1h at 4 °C under gentle agitation in the presence of 1 mg/ml BSA. Beads were washed 5x with a 50-fold excess of binding buffer without BSA and mixed with SDS-sample buffer. Samples were boiled and an equivalent of 5 µl of beads was analyzed by SDS PAGE. Stained gels were scanned, and band intensity was quantified by measuring the mean gray value in FIJI.

### Co-filtration of nanodiscs with Osh6

The Osh6 mutant C389S was purified form *S. cerevisiae* as described for WT and the remaining surface accessible cysteine (Cys 62) was labeled with the fluorophore Cy5 using the Protein labeling kit ‘RED-MALEIMIDE 2nd Generation’ (Nanotemper). Labeled Osh6 was incubated with a 1.2-fold excess of nanodiscs composed of a MSP 2N2/POPG:POPC (3:1) mixture at a molar ratio of 1:200 in and injected on a Superose 6 5/150 GL column. The eluate was collected in 100 µl fractions and the Cy5 fluorescence of the fractions was measured in a TECAN Sparc plate reader.

### Isothermal titration calorimetry

ITC measurements were performed in a MicroCal iTC200 calorimeter at 25 °C with an initial delay of 60 s and stirring at 1000 rpm. Ist2 peptides at a stock concentration of 240 µM were titrated with 20 injections of 2 µl aliquots (except the first injection containing 1.4 µl) with a spacing of 180 s, reference power of 3DP and a filter period of 5 s at high feedback mode/gain. Osh6 was filled in the reaction cell aiming at a concentration of 20 µM, the exact concentration used for data fitting was determined for each measurement from superfluous liquid after filling the reaction cell. The reference cell contained H_2_O. Osh6 and the Ist2^732–761^ peptide were in the same buffer (25 mM HEPES pH 7.5, 150 mM NaCl, 5% glycerol). The baseline was subtracted by titrating the analyte into buffer. Data was fitted in Origin using a single site sigmoidal fit in an iterative manner until the chi^2^ values converged.

### X-Ray crystallography

Peptides used for crystallography and ITC (Ist2 732-761 and Ist2 732-757) were purchased from Genscript. Osh6^Δ30^ was concentrated to 200 µM after affinity purification and cleaved overnight upon addition of 3C protease. The cleaved protein was incubated for 30 min with a 1.2 molar excess of the respective peptide and injected onto a Superdex 200 10/300 GL column equilibrated in 15 mM HEPES pH 7.8, 150 mM NaCl. Osh6 peak fractions were pooled and concentrated to 18.5 mg/ml. The presence of the peptide was confirmed by mass-spectroscopy. Crystallization experiments were performed at the protein crystallization facility of the Dept. of Biochemistry of UZH. The final sample was supplemented with an additional 0.5 molar excess of peptide and mixed with the crystallization liquor at a ratio of 1:1 at 20 °C. Osh6/Ist^732–757^ crystallized at 20 °C in 100 mM BisTris pH 5.5, 200 mM KCl, 75 mM MgCl_2_, 28-29% PEG 400. Osh6/Ist^732–761^ crystallized at 4 °C in 100 mM BisTris pH 5.5, 200 mM KCl, 20 mM MgCl_2_, 28%-30% PEG400. Crystals formed within 10 days and were cryoprotected by stepwise increase of the PEG400 concentration to 35% before flash freezing in liquid nitrogen. X-ray diffraction data was collected at the Petra III p13 beamline at EMBL Hamburg. Data was processed with XDS^56^ and EDNA^57^. The crystals were of space group C222_1_ with one copy of the complex in the asymmetric unit and Data extended to 1.9 Å for both datasets (Table S2). The structure was solved using molecular replacement with Phaser^58^ implemented in the Phenix software package^59^ using the coordinates of Osh6 (PDB ID 4B2Z) as start model. Iterative refinement and model building was performed in Coot^60^ and Phenix^61^.

### Sample preparation for cryo-EM and data collection

For Ist2^716^ in GDN, the protein was concentrated to 2.56 mg ml^-1^ and 2.5 µl of sample was applied to a Quantifoil R1.2/R1.3 Au200 grid, blotted for 2 s and plunge frozen in a liquid ethane/propane mix using a Mark V Vitrobot. Data was collected on a Titan Krios G3i equipped with a post column energy filter and a Bioquantum K3 detector at a physical pixel size of 0.651 Å at a magnification of 130,000. Further samples were prepared similarly with the following exceptions: Ist2 1-716/MSPE1D3 was concentrated to 1.09 mg ml^-1^ and applied to a Quantifoil UltrAuFoil R1.2/1.2 300 grid. Osh6/MSP2N2 was concentrated to 1.63 mg ml^-1^ and blotted for 3 s before plunging.

### Cryo-EM data collection and processing

All raw movies were gain, motion and CTF corrected in cryoSPARC version 4 (ref. ^62^). Micrographs with poor CTF estimation, high drift and ice thickness were discarded. For the Ist2/Osh6 dataset, the blob picker function was used to generate Templates from a random subset of 1000 micrographs, which represented different particle sizes and the heterogeneity of the dataset and were subsequently used for Template picking. After an initial 2D classification of all template particle picks, the particles were sorted into three subsets based on their size and the number of particles in the box. Classes exhibiting features of the Ist2 transmembrane domain were subjected to multi-class ab-initio reconstruction yielding a single class with well-defined transmembrane density. Particles sorted into this class were used to train a Topaz model to increase the number of picks^63^. After re-extraction the picked particles and further cleaning of the particle set using 2D classification, the particles were again subjected to ab-initio reconstruction and the single good class was further refined using heterogenous refinement with a decoy class and homogenous refinement in C1. Since the map exhibited strong C2 features, a final round of non-uniform refinement with imposed C2 symmetry was performed to obtain an improved final reconstruction. Classes with small particles were subjected to separate ab-initio reconstruction, resulting in two classes with triangular shape which in shape and size resembled previously solved crystal structures of Osh6. Since pooling of those classes did not improve the result, the best class was chosen for homogenous refinement with a large, soft mask, resulting in an envelope of the 50 kDa protein at 11 Å resolution. For the Ist2^716^ structure in GDN, templates for template picking were generated from a subset of 100 micrographs using the blob picker function. Template picks were subjected to several round of 2D classification until the resolution of classes did not improve further and particles were no longer sorted into junk classes. The clean particle set was subjected to ab-initio reconstruction with two classes, resulting in a single good ab-initio model, which was refined using heterogenous refinement with a decoy class generated from the same dataset and subjected to non-uniform refinement in C1. Upon reaching Nyquist, particles contributing to the final reconstruction were unbinned and refined with global CTF refinement and C2 symmetry imposed to obtain the final reconstruction. For the Ist2^716^ structure in 1E3 nanodiscs, templates were generated from a subset of micrographs using blob picker. The initial particle set was cleaned by using iterative rounds of 2D classification. Particles sorted into 2D classes with visible transmembrane density were unbinned through re-extraction and after further 2D classification subjected to multi-class ab-initio reconstruction, resulting in two classes with well-defined transmembrane density differing in the apparent size of the nanodisc as well as the visibility of the cytosolic bridging helix and features of the lower half of α6. The two distinct classes and two junk classes obtained from ab-initio reconstruction were used as input models for sorting of all particles using heterogenous refinement and then refined separately using non-uniform refinement. In case of the “narrow” conformation, C2 symmetry improved the resolution and was therefore applied in the final refinement step. In cases of the Ist2fl/Osh6 complex and Ist2^716^_ND_ structures, both local and global CTF refinement did not improve the maps. For the Osh6 Nanodisc Dataset, initial particle picking was performed using blob picker in cryoSPARC. The particle picking was then improved in sequential rounds of 2D classification and Topaz Picking. Particles were subjected to ab-initio classification, which resulted in classes showing free nanodiscs and Nandiscs bound to a molecule corresponding to approximate size of Osh6. The particle set was further sorted in 3D using heterogenous refinement to exclude unbound Nanodiscs and subjected to an initial refinement in cryoSPARC. The refinened particles were then exported and re-extracted in Relion v5 with a smaller box size. The particles were then subjected to 2 iterative rounds of unmasked 3D classification without alignment. Particles from classes showing higher resolution features of Osh6 were re-extracted with re-centering on Osh6 and subjected to another round of 3D classification without alignment with a soft mask including the Osh6 density. The two best classes were pooled and aligned using the auto-refine option of Relion v5^64,65^. The density corresponding to the Nanodisc was subtracted by masking the Osh6 density. The particles were subjected to a final refinement and Relion postprocessing. Details of the data processing workflow are shown in Extended Data Figures.

### Yeast growth assays

Ist2 constructs were transformed into a *S. cerevisiae ist2/psd1* KO strain (AWY327), generously provided by Dr. Chris Loewen, using the LiAc/SS carrier DNA/PEG method^54^ and plated onto SC-Ura selection medium containing 1mM Choline Bitartrate. Plates were kept at 4 °C for maximally 4 weeks. Single colonies were picked and grown in liquid SC-Ura medium containing 1 mM Choline to mid-log phase. Cells were harvested by centrifugation and pellet was washed 3 times with sterile water to remove all traces of Choline. Cells were subsequently diluted to OD 0.35 and a 10x dilution series was spotted in three replicates on solid media containing 2% dextrose and 2% galactose or 1mM choline. Plates were incubated at 30 °C for 3-4 days. The experiment was repeated 3 times with different transformants for each construct. For evaluation, plates were imaged in a Vilber Fusion FX using bright field illumination. The background was subtracted in FIJI and the mean grey value of each spot was quantified using a circular mask of equal size for all replicates and all constructs. The quantification of the 1:100 dilution was chosen for comparison between constructs due to being in the linear range of the 1:10 dilution series. For each replicate, all constructs were normalized to Ist2 WT. Statistical analysis was performed in GraphPad Prism using ANOVA.

## Data availability

Cryo-EM density maps and atomic models will be deposited in the Protein Data Bank and Electron Microscopy Data Bank and will be publicly available as of the date of publication. The PDB IDs are 9HDH (Osh6ΔN30^732–761^), 9HDK (Osh6ΔN30^732–756^), 9HHE (Ist2^716^). EMDB IDs are EMD-52170 (Ist2^716^_GDN_). For review, the coordinates and structure factors for X-ray datasets and coordinates maps and half-maps for cryo-EM datasets are deposited with the manuscript.

---

**Extended Data Fig. 1.**
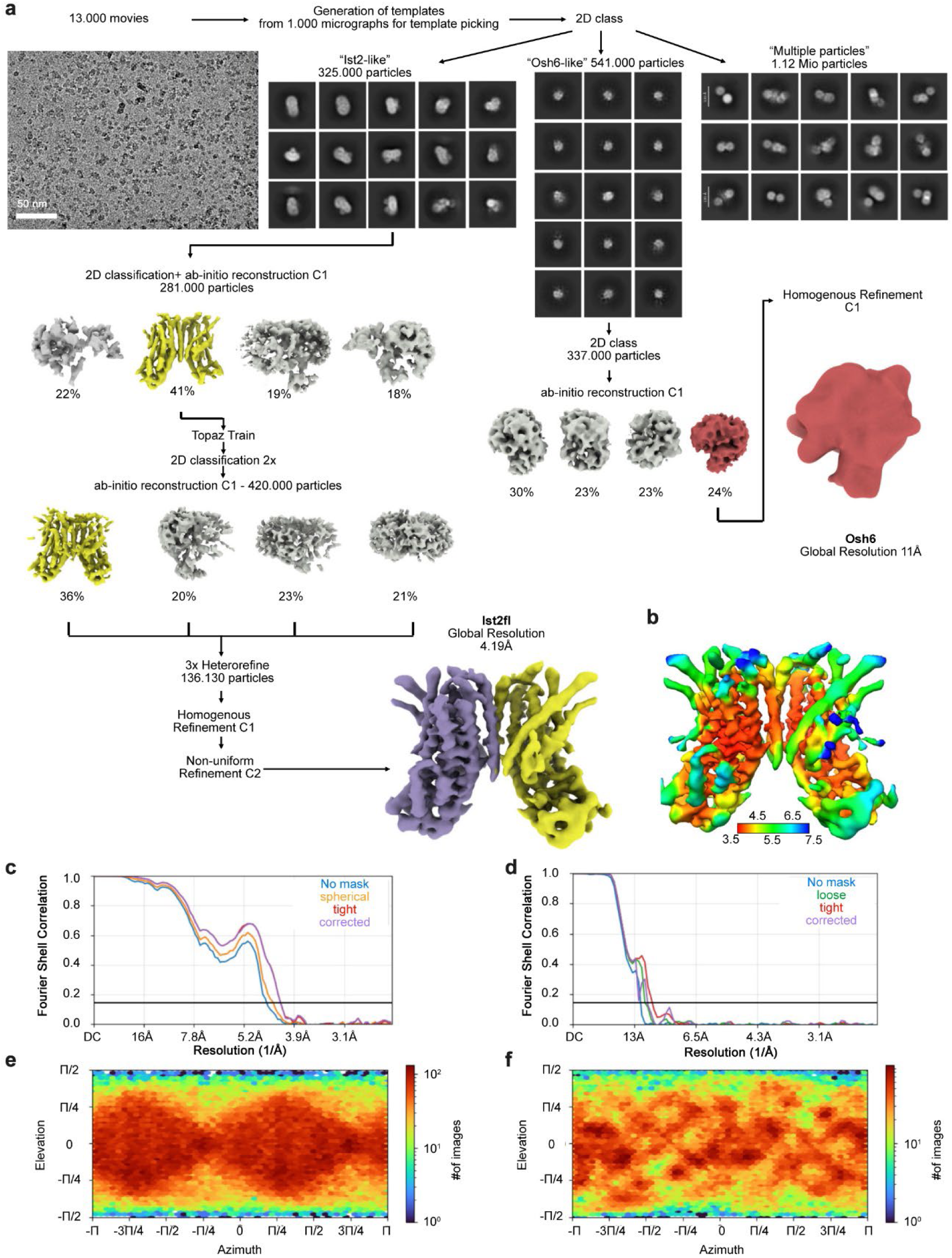
Cryo-EM data processing of Ist2 in detergent in complex with Osh6. **a**, Representative micrograph, 2D classes and data processing workflow for Ist2/Osh6 complex. Scalebar on micrograph is 50 nm. Particles were sorted in 2D into three categories based on size. Ist2 and Osh6 particles were then processed separately as indicated. Data was processed using cryoSPARC. **b**, Local resolution of the final Ist2 map computed in cryoSPARC. **c**, GSFSC curve used for resolution estimation of final maps of Ist2. **d**, GSFSC curve used for resolution estimation of final maps of Osh6. **e**, Angular viewing direction distribution of particles contributing to the final map of Ist2fl. **f**, Angular viewing direction distribution of particles contributing to the final map of Osh6.

**Extended Data Fig. 2.**
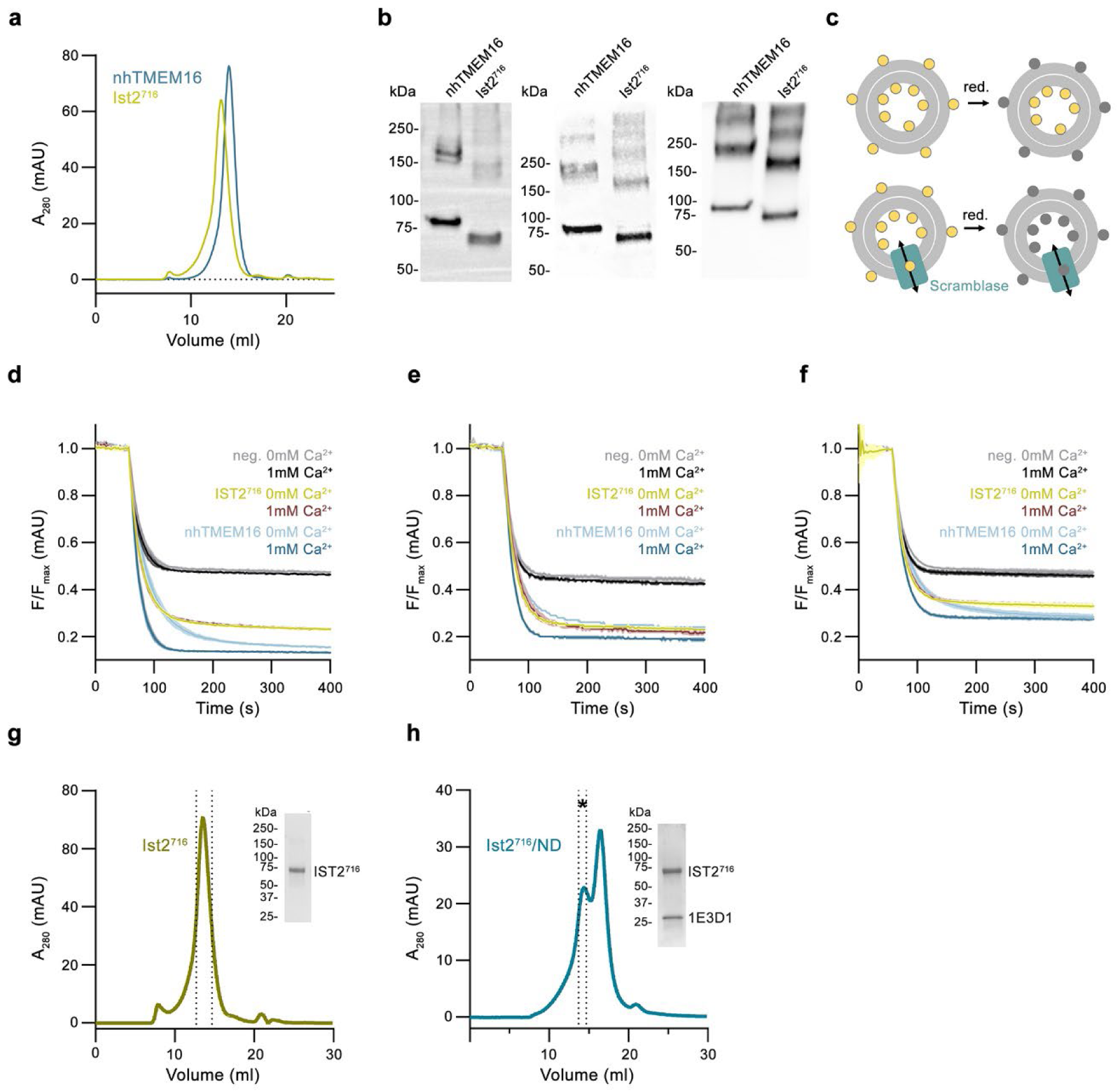
Biochemical properties of Ist2^716^ and lipid transport experiments. **a**, Size exclusion chromatogram of a sample of Ist2^7^^16^ and nhTMEM16 used for liposome reconstitution. **b**, SDS Page gels of proteoliposomes containing nhTMEM16 and Ist2^7^^16^ from three independent reconstitutions used for scrambling assays. **c**, Scheme of the lipid scrambling assay. ‘red.’ refers to the addition of dithionite to the outside of liposomes. Fluorescent lipids are indicated as yellow, lipids with bleached fluorophore as grey spheres, a scramblase mediating the bidirectional diffusion of lipids between leaflets as cyan box. **d**-**f**, Scrambling data from three independent reconstitutions that are combined in Fig. 1f, g. The displayed nhTMEM16 and Ist2^7^^16^ preparations were reconstituted into the same batch of destabilized liposomes. Empty liposomes of the same batch are shown as negative control (neg.). Recordings show mean of three technical replicates, errors are s.d. **g**-**h**, Size exclusion chromatograms of Ist2^7^^16^ in detergent (**g**) and after reconstitution in lipid nanodiscs (**h)** used as samples for the cryo-EM datasets shown in Extended Data Figs. 3 and 4, respectively. Dashed lines indicate pooled fractions.

**Extended Data Fig. 3.**
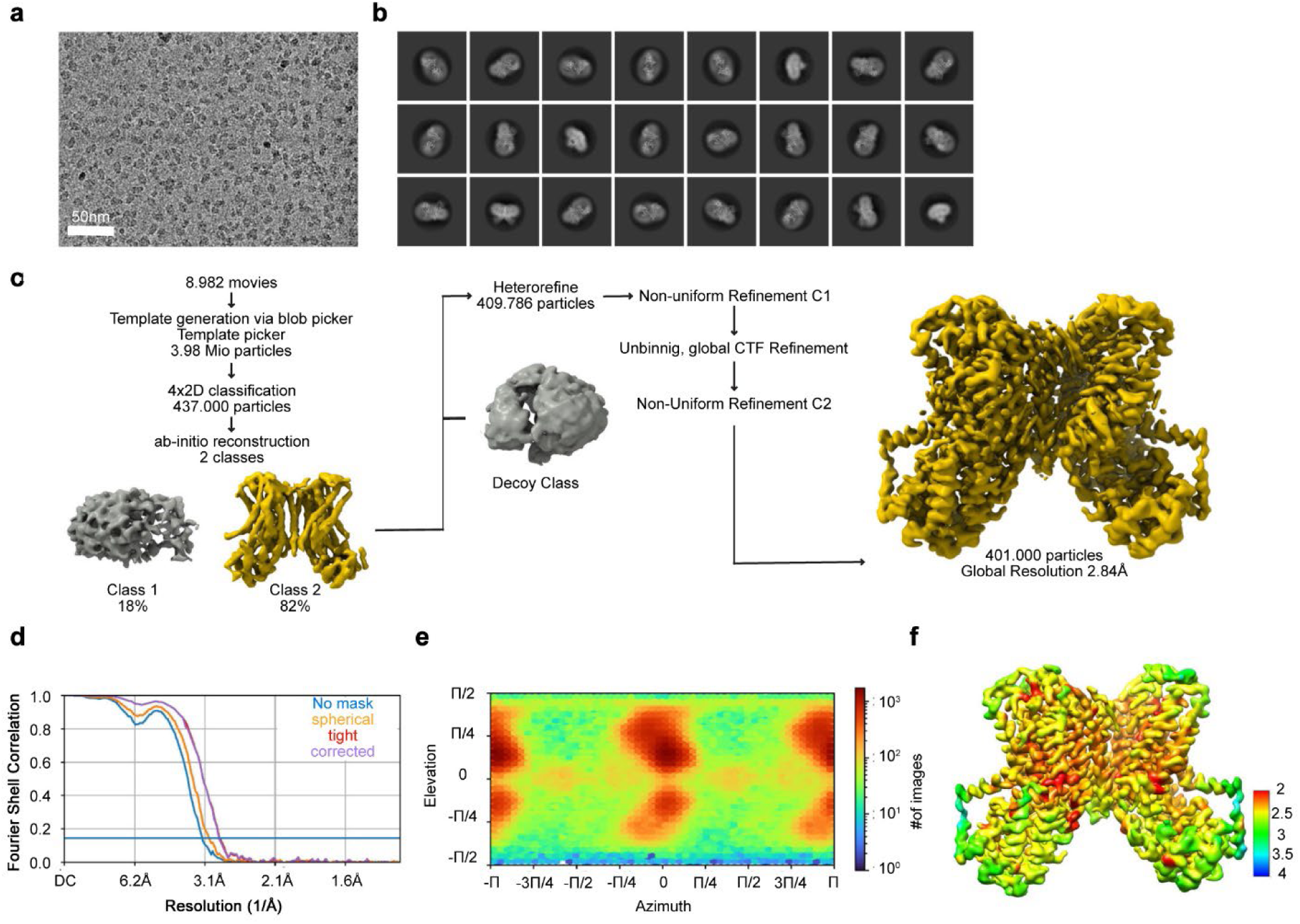
Cryo-EM data processing of the Ist2 transmembrane domain in detergent. **a**, Representative CTF-corrected micrograph for Ist2^716^_GDN_, scale bar is 50 nm. **b**, Representative final 2D classes for Ist2^716^_GDN._ **c**, Data processing workflow for Ist2^716^_GDN_ performed in cryoSPARC. **d,** GSFSC curve used for resolution estimation of final map of Ist2^716^_GDN_. **e**, Angular viewing direction distribution of particles contributing to the final map of Ist2^716^_GDN_. **f**, Local resolution estimation for Ist2^716^_GDN_ computed in cryoSPARC.

**Extended Data Fig. 4.**
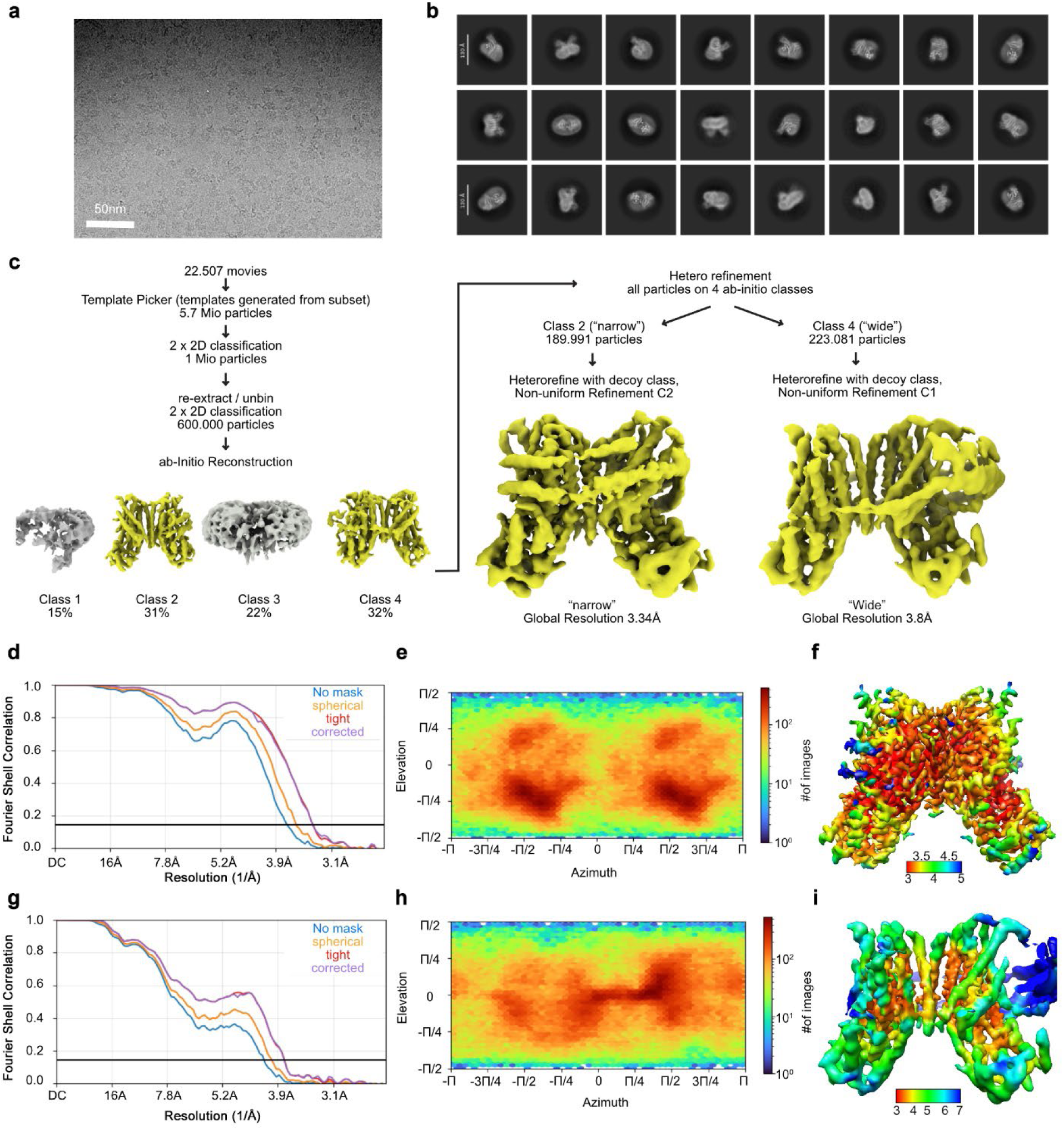
Cryo-EM data processing of the Ist2 transmembrane domain in lipid nanodiscs. **a**, Representative CTF-corrected micrograph for Ist2^716^_ND_, scale bar is 50 nm. **b**, Representative final 2D classes for IST Ist2^716^_ND._ **c**, Data processing workflow for Ist2^716^_ND_ performed in cryoSPARC leading to two distinct reconstructions based on nanodisc size, termed “narrow” and “wide”. **d**, GSFSC curve used for resolution estimation of final map of Ist2^716^_ND_ “narrow” conformation. **e**, Angular viewing direction distribution of particles contributing to the final map of Ist2^7^^16^ Ist2^716^_ND_ “narrow”. **f**, Local resolution estimation for Ist2^716^_ND_ “narrow” computed in cryoSPARC. **g**, GSFSC curve used for resolution estimation of final map of Ist2^716^_ND_ “wide” conformation. **h**, Angular viewing direction distribution of particles contributing to the final map of Ist2^7^^16^ Ist2^716^_ND_ “wide”. **i**, Local resolution estimation for Ist2^716^_ND_ “wide” computed in cryoSPARC.

**Extended Data Fig. 5.**
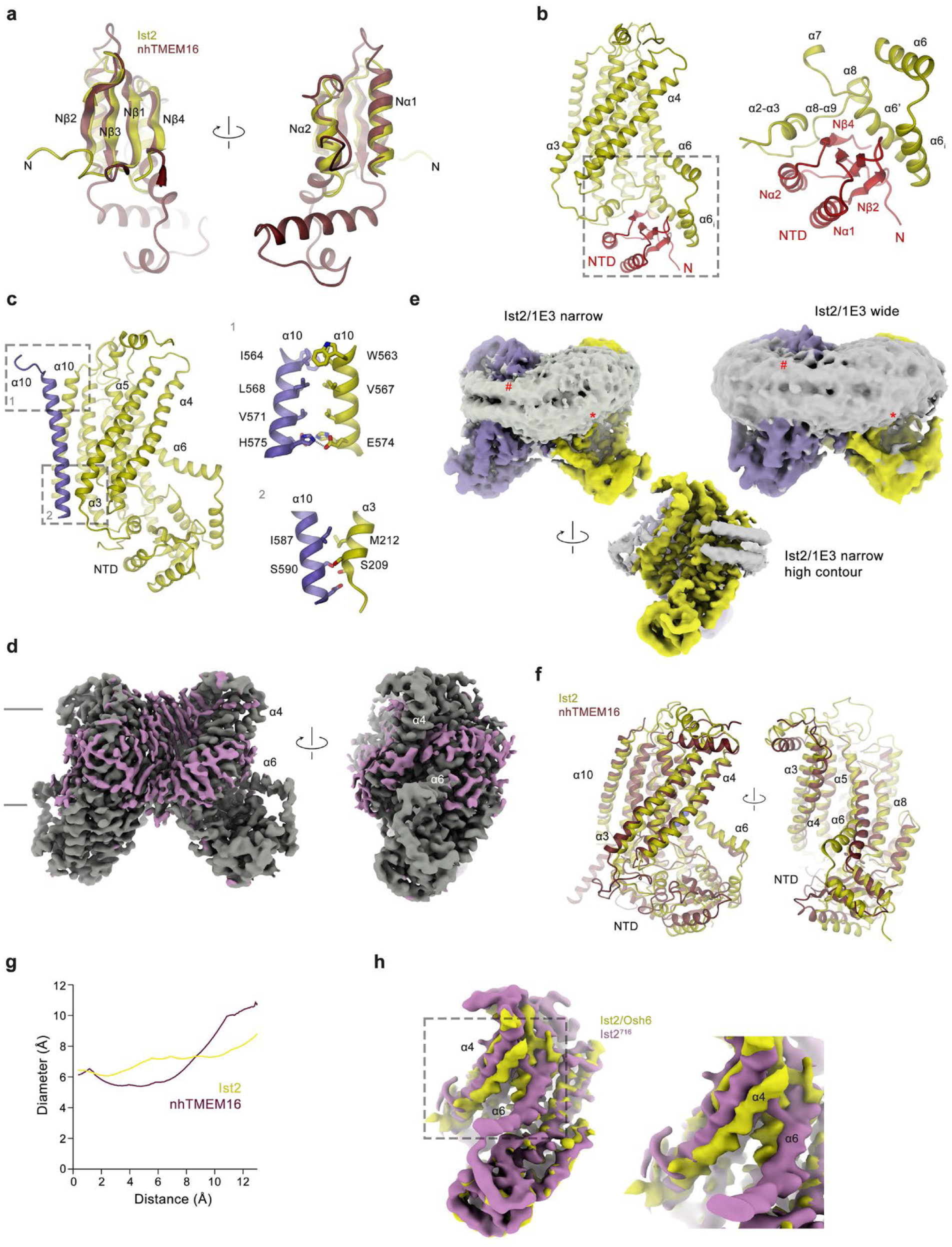
Features of the Ist2 transmembrane domain. **a**, Superposition of NTDs of Ist2 and nhTMEM16. Secondary structure elements of Ist2 are labeled. **b**, Location of the NTD (red) in the Ist2 subunit. Ribbon representation of the subunit is shown left. Blowup of the marked region right. **c**, Dimer interface. Left, Ist2 subunit with α10 of the contacted subunit displayed as ribbon. Boxes indicate magnified regions displayed right. Top, interacting residues between symmetry related helices α10 and bottom, the α10-α3 interface. Sidechains of interacting residues are shown as sticks. **d**, Cryo-EM densities of Ist2^7^^16^ constructs in nanodiscs showing narrow (left) and wide (right) particle reconstructions. A view of the narrow reconstruction along the dimer (bottom) at larger contour illustrates presumable density of the scaffolding protein. Membrane deformations are indicated (#, *). **e**, Bulky lipid-like density (magenta) surrounding the membrane-inserted region of Ist2^7^^16^ detergent dataset. **f**, Superposition of subunits of the ‘open’ conformations Ist2 and nhTMEM16 (PDBID 6QM9). **g**, Diameter of the open subunit cavity in Ist2 and nhTMEM16 measured by Hole. **h**, Superposition of the densities of the membrane inserted part of the Ist2/Osh6 complex and the Ist2^7^^16^ subunit (low pass filtered to the same resolution) illustrates a different conformation of α4 and α6. A blowup of the marked region is shown right.

**Extended Data Fig. 6.**
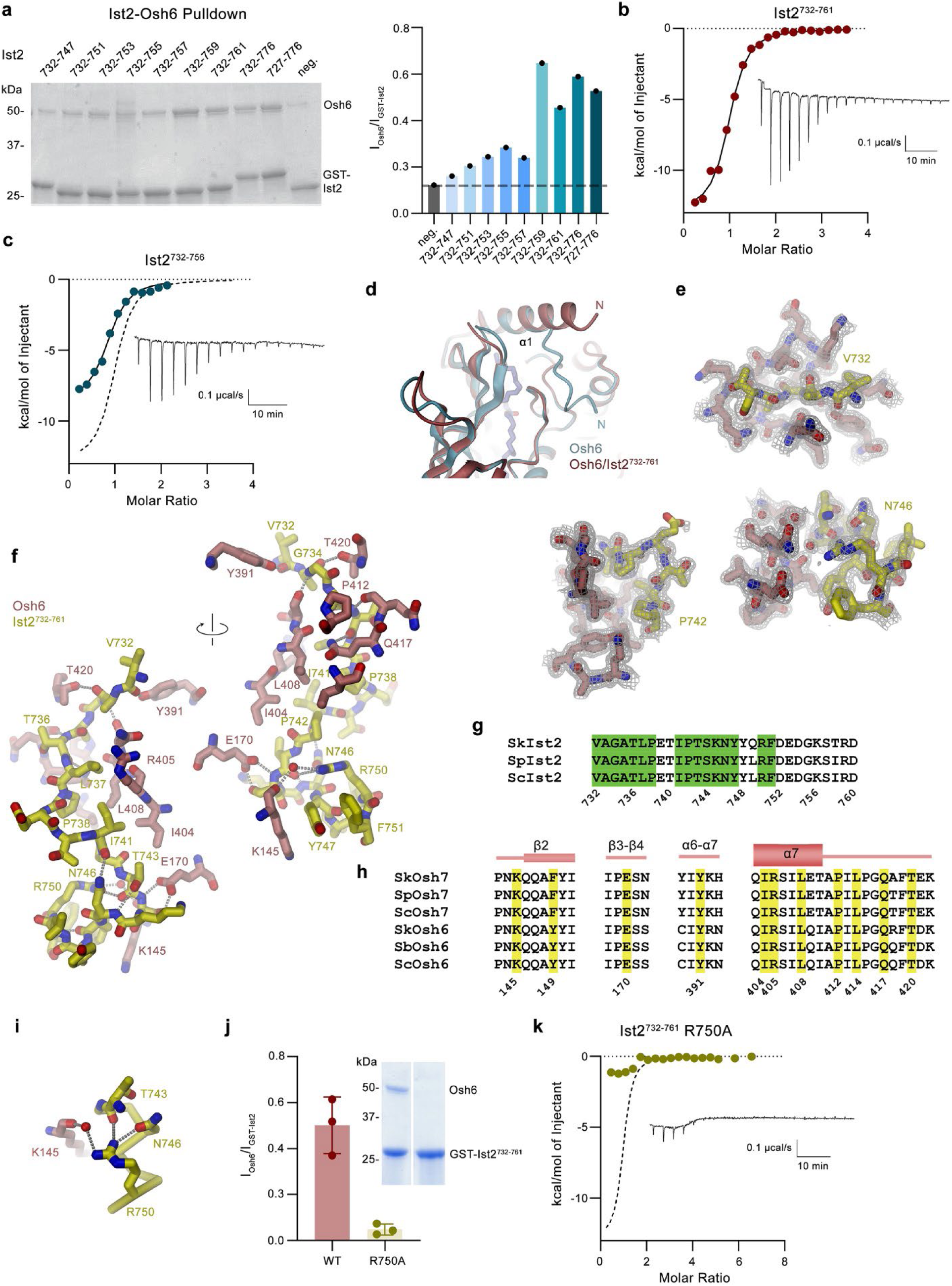
Biochemical and structural characterization of the Ist2/Osh6 interaction. **a**, Left, SDS-PAGE gel of pulldown of Osh6 with fusion-constructs of peptides carrying the indicated region of Ist2 fused to GST by binding to GSH resin. Neg. refers to GST not containing any attached peptide. Intensity ratio of bands corresponding to Osh6 and the respective GST-Ist2 fusion constructs. Dashed line indicates value of negative control (neg.). Values were obtained from a single experiment. **b**, **c**, Binding of a peptide encompassing residues 732-761 of Ist2 (Ist2^732–761^, showing an independent biological replicate from data displayed in Fig. 3a) (**b**) or residues 732-756 (Ist2^732–756^) (**c**) to Osh6. Binding isotherms were obtained from isothermal titrations of either peptide to Osh6. The data show fits to a model assuming a single binding site with the binding isotherm depicted as solid line with K_d_ of 714.0 ± 86.7 nM (Ist2^732–761^) and a K_d_ of 1.1 ± 0.17 μM (Ist2^732–756^). The dashed line in b refers to the fit of Ist2^732–761^. Errors represent fitting errors. The thermograms are shown as inset (right). **d,** Superposition of Osh6 structures obtained in absence (cyan, PDBID 2B2Z) and in complex with Ist2^732–761^ (red). The displayed region illustrates conformational differences in the lid helix α1 covering the lipid binding site. **e**, 2Fo-Fc density (grey mesh) superimposed on different regions of the Osh6/Ist2 interface. **f**, Structural details of Ist2-Osh6 interactions. **g**, Sequence alignment of Osh6 binding region of Ist2 of three Saccheromyces species (*S. paradoxus, S. kudriavzeii, S. cerevisiae*). Residues in contact with Osh6 are colored in green. Numbering refers to the *S. cerevisiae* Ist2. **h**, Sequence alignment of Ist2 interactions of Osh6 and Osh7 of the same strains. Residues in contact with Ist2 are colored in yellow. Numbering refers to the *S. cerevisiae* Osh6. **i**, Intra and intermolecular Interactions of the Arg 750 sidechain of Ist2. **j**, Pulldown of Osh6 by a peptide of the Ist2^732–761^ region carrying the point mutation R750A in comparison to WT. Experiments were caried out as described for A. Bars show mean of three technical replicates, errors are s.d.. **k**, Binding of a peptide encompassing residues 732-761 of Ist2 carrying the mutation R750A to Osh6. Binding isotherms were obtained from isothermal titrations of the peptide to Osh6. Dashed line refers to the fit of Ist2^732–761^. The thermogram is shown as inset (right).

**Extended Data Fig. 7.**
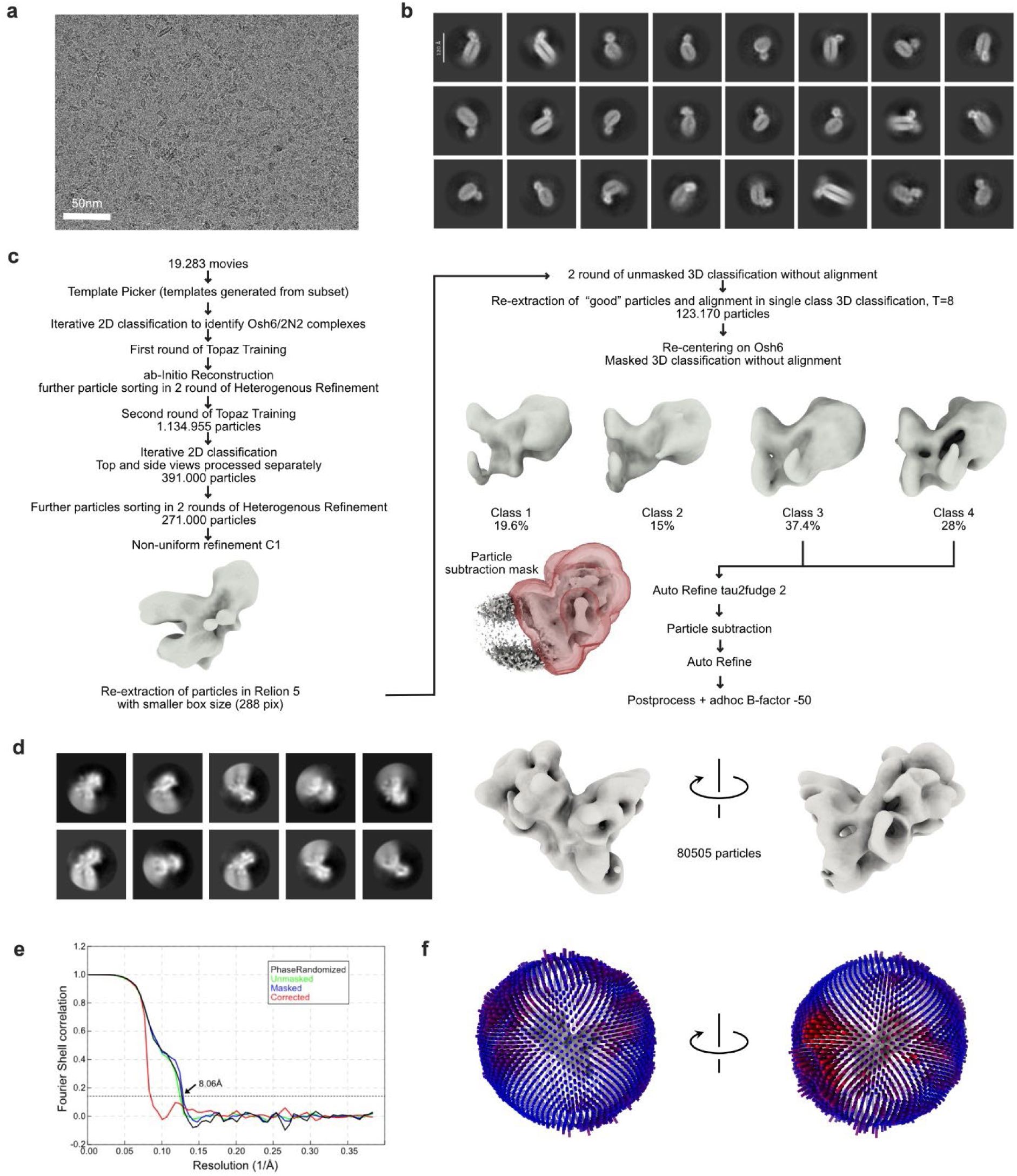
Structural characterization of the Osh6 interaction with lipid nanodiscs. **a**, Representative CTF-corrected micrograph for Osh6_ND_, scale bar is 50 nm. **b**, 2D classes for Osh6_ND_ calculated in cryoSPARC representing the Osh6-bound fraction of Nanodiscs used for initial 3D reconstruction. **c**, Data processing workflow for Osh6_ND_ performed in cryoSPARC and Relion v5. **d**, Result of 2D classification without alignment of refined particles in the final map performed in Relion v5. **e**, GSFSC curve of final map displayed in (c) computed in Relion v5. **f**, Euler angle distribution for final 3D reconstruction shown for two orientations of the map.

**Extended Data Fig. 8.**
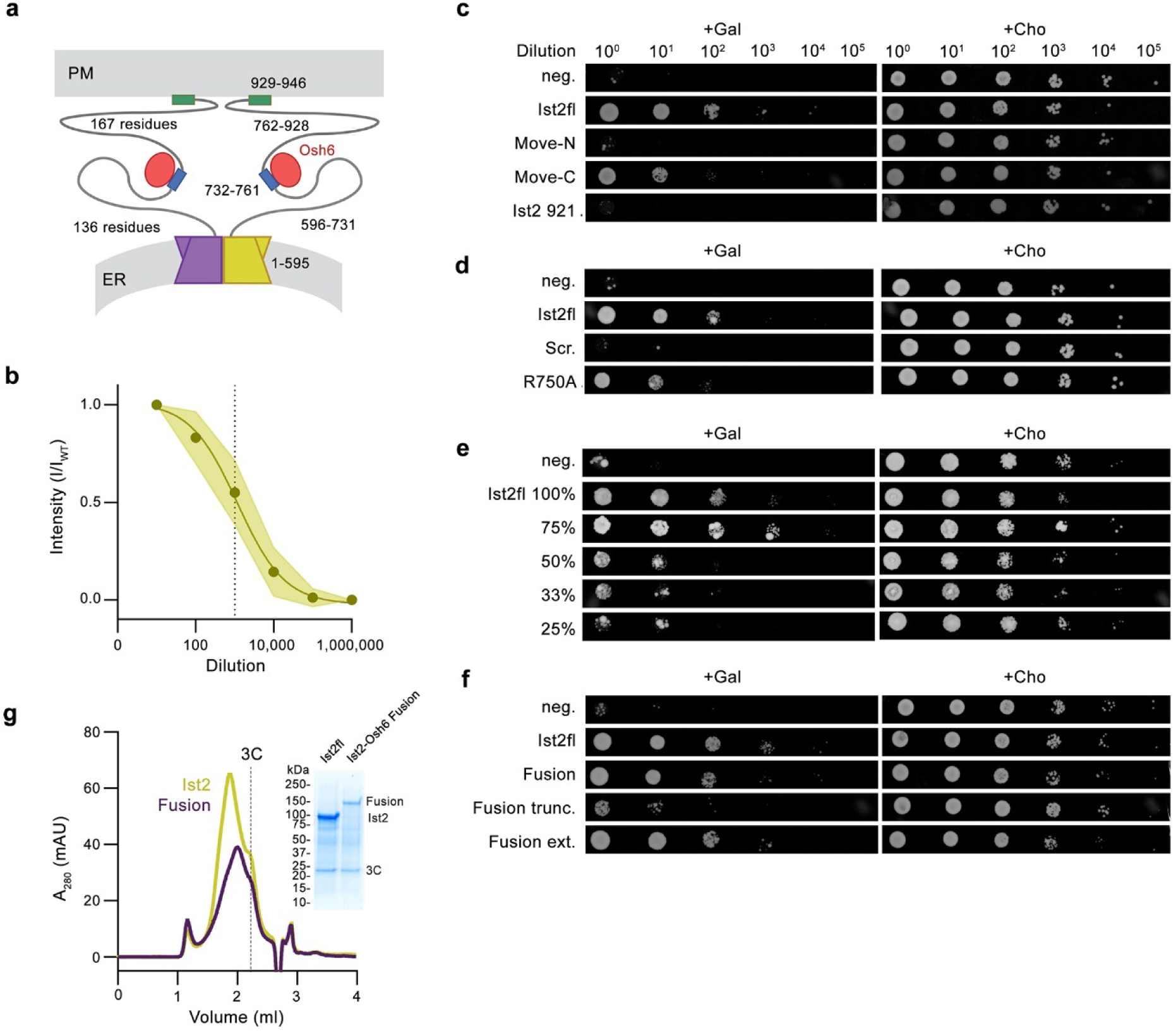
Functional characterization of Ist2 constructs by yeast complementation assays, and fluorescence microscopy. **a**, Scheme of the Ist2/Osh6 complex bridging the ER and the PM. The residue range of the indicated parts of Ist2 is shown. **b**, Normalized mean gray value quantification of all 10-fold spot assay dilution series of Ist2 transformed into *ist2/psd1* KO cells under inducing conditions (2% Galactose, n=27). Dashed line indicates 1:100 dilution, which was used for evaluation and comparison of all other constructs. For each dilution series, values were normalized to the maximum and minimum mean gray value. Data was fit with a sigmoidal model using GraphPad prism. **c**-**f**, Yeast spot dilution series used for FIJI-based evaluation shown in Fig.6. Shown are **c**, WT in comparison to the construct Ist2^9^^21^ lacking the C-terminal attachment site to the PM and constructs where the Osh binding site was moved by 83 residues in N or C-terminal direction of the linker, **d**, mutations in the Osh6 binding region, **e**, constructs with decreasing liker length, **f**, fusions of Osh6 into the linker. Images show representative 10-fold dilution series of n=3x3 (n=2x3 for Fusion trunc. and Fusion ext.) replicates for plates containing 2% galactose and 1mM choline for each Ist2 construct transformed into *ist2/psd1* KO cells. Neg. refers to cells transfected with pYESnSM3 vector only. Plates were imaged with a Vilber Fusion FX detection system and the background was subtracted using FIJI. **g**, Size exclusion chromatogram of the purified Ist2-Osh6 fusion construct (Fusion) in comparison to Ist2 WT.

**Extended Data Table 1.** Summary of EM Data and refinement statistics.

|  | #1 Ist2<sup>716</sup><sub>GDN</sub><br>(EMDB-52170)<br>(PDB 9HHE) | #2 Ist2<sup>716</sup><sub>ND</sub><br>narrow | #3 Ist2<sup>716</sup><sub>ND</sub><br>wide | #4 Ist2<sup>716</sup><sub>fl</sub> |
| --- | --- | --- | --- | --- |
| **Data collection and processing** |  |  |  |  |
| Magnification | 130.000x | 130.000x | 130.000x | 130.000x |
| Voltage (kV) | 300 | 300 | 300 | 300 |
| Electron exposure (e<sup>-</sup>/Å<sup>2</sup>) | 62.43 | 62.43 | 62.43 | 62.43 |
| Defocus range (μm) | -1 to -2.4 | -1 to -2.4 | -1 to -2.4 | -1 to -2.4 |
| Pixel size (Å) | 0.651 | 0.651 | 0.651 | 0.651 |
| Symmetry imposed | C2 | C2 | C1 | C2 |
| Initial particle images (no.) | 3.981.000 | 5.7000.000 | 5.7000.000 | 420.000 |
| Final particle images (no.) | 400.925 | 189.991 | 223.081 | 136.130 |
| Map resolution (Å) | 2.84 | 3.34 | 3.8 | 4.19 |
| FSC threshold 0.143 |  |  |  |  |
| Map resolution range (Å) | 2-3.5 | 3-4.5 | 3.5-6 | 3.5-6.5 |
| **Refinement** |  |  |  |  |
| Initial model used (PDB code) | Ist2 AlphaFold | - | - | - |
| Model resolution (Å) | 2.8 |  |  |  |
| FSC threshold 0.5 |  |  |  |  |
| Map sharpening B factor (Å<sup>2</sup>) | -50 |  |  |  |
| Model composition |  |  |  |  |
| Non-hydrogen atoms | 9248 |  |  |  |
| Protein residues | 1130 |  |  |  |
| Ligands | - |  |  |  |
| B factors (Å<sup>2</sup>) |  |  |  |  |
| Protein | 85.57 |  |  |  |
| Ligand | - |  |  |  |
| R.m.s. deviations |  |  |  |  |
| Bond lengths (Å) | 0.002 |  |  |  |
| Bond angles (°) | 0.549 |  |  |  |
| Validation |  |  |  |  |
| MolProbity score | 1.93 |  |  |  |
| Clashscore | 7.58 |  |  |  |
| Poor rotamers (%) | 2.01 |  |  |  |
| Ramachandran plot |  |  |  |  |
| Favored (%) | 95.87 |  |  |  |
| Allowed (%) | 3.95 |  |  |  |
| Disallowed (%) | 0.18 |  |  |  |

**Extended Data Table 2.** Data collection and refinement statistics of X-ray structures of Osh6 complexes.

|  | Osh6ΔN30<sup>732-761</sup><br>(PDB 9HDH) | Osh6ΔN30<sup>732-756</sup><br>(PDB 9HDK) |
| --- | --- | --- |
| **Data collection** |  |  |
| Space group | C222<sub>1</sub> | C222<sub>1</sub> |
| Cell dimensions |  |  |
| *a*, *b*, *c* (Å) | 61.64, 176.25, 95.71 | 61.37, 175.7, 96.16 |
| $\alpha$, $\beta$, $\gamma$ (°) | 90, 90, 90 | 90, 90, 90 |
| Resolution (Å) | 47.86 - 1.84 | 50.0 - 1.93 |
| *R*<sub>merge</sub> | 19.6 (154.1)* | 8.3 (149.3)* |
| *I* / σ*I* | 7.91 (1.38) | 18.0 (1.8) |
| Completeness (%) | 97.9 (87.1) | 99.6 (99.4) |
| Redundancy | 6.73 (5.19) | 13.7 (14.0) |
| **Refinement** |  |  |
| Resolution (Å) | 1.84 | 1.93 |
| No. reflections | 44425 | 41459 |
| *R*<sub>work</sub> / *R*<sub>free</sub> | 17.95/20.97 | 18.51/21.32 |
| No. atoms |  |  |
| Protein | 3485 | 3422 |
| Ligand/ion | 53 | 53 |
| Water | 319 | 315 |
| *B*-factors |  |  |
| Protein | 31.79 | 43.84 |
| Ligand/ion | 32.02 | 39.03 |
| Water | 38.13 | 49.21 |
| R.m.s. deviations |  |  |
| Bond lengths (Å) | 0.0067 | 0.0064 |
| Bond angles (°) | 0.839 | 0.822 |

\* Values in parenthesis refer to highest resolution shell

**Supplementary Figure 1.**
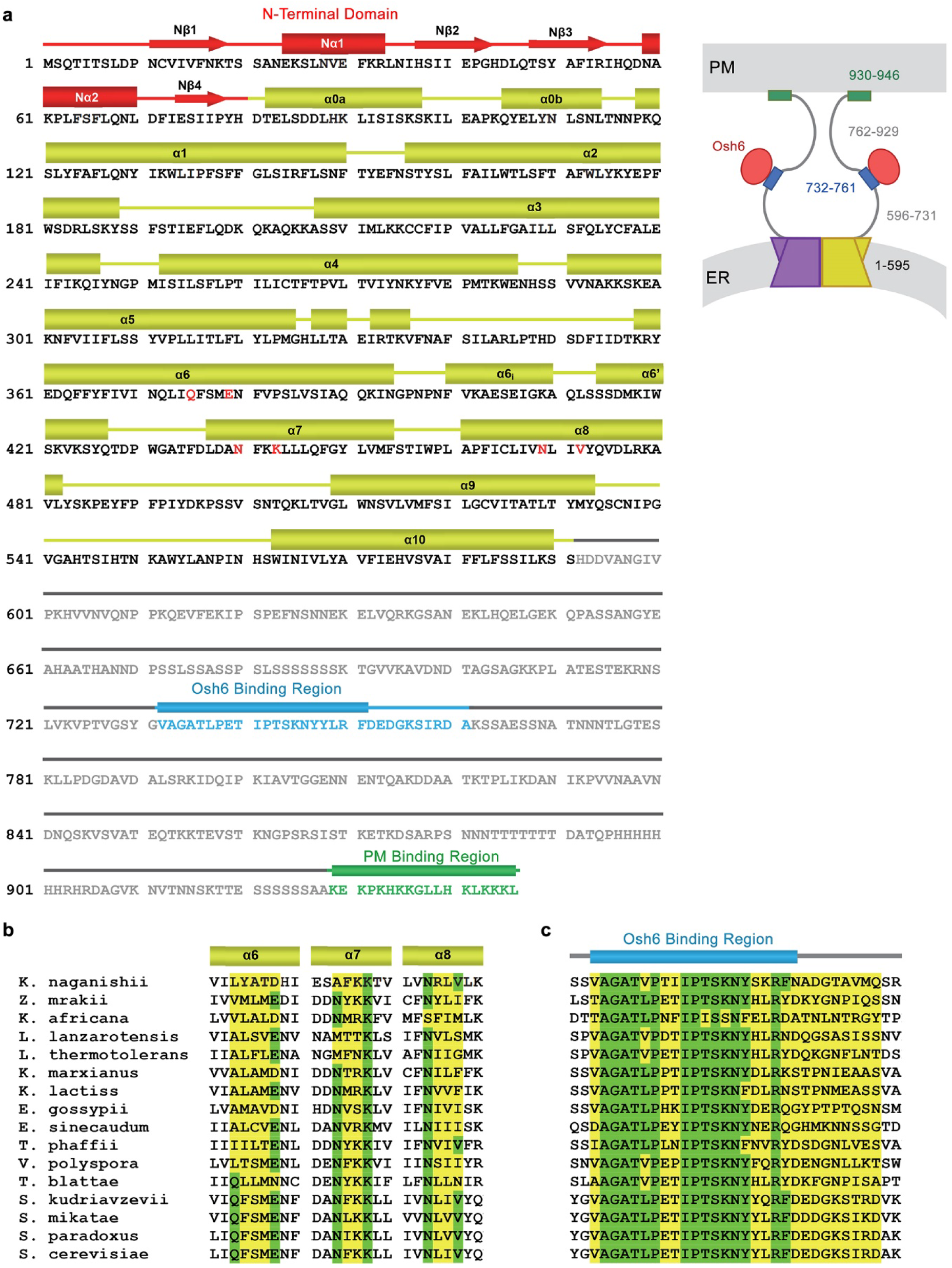
Ist2 sequence. **a**, Sequence of Ist2 from *S. cerevisiae*. Secondary structure elements and regions involved in Osh6 and PM binding are indicated above. Structured regions are shown in black, unstructured regions in grey, residues of the Osh6 binding site in blue and residues interacting with the PM in green. A scheme of the Ist2/Osh6 complex bridging the ER and the PM is shown right. The residue range of the different parts of Ist2 is indicated. **b**, Sequence alignment of the region defining the regulatory Ca^2+^ binding site in the TMEM16 family in Ist2 homologs from *Saccharomycetaceae*. Secondary structure elements are shown above. Residues equivalent to the binding region are highlighted in yellow, conserved residues in green. **c**, Sequence alignment of the region defining the regulatory Osh6 binding site in Ist2 homologs from *Saccharomycetaceae*. The location of residues in direct interaction with Osh6 is indicated. The wider region is colored in yellow, identical residues in green. **b**, **c**, Sequences are of Ist2 homologs from *Kazachstania naganishii, Zygotorulaspora mrakii, Kazachstania africana, Lachancea lanzarotensis, Lachancea thermotolerans, Kluyveromyces marxianus, Kluyveromyces lactis, Eremothecium gossypii, Eremothecium sinecaudum, Tetrapisispora phaffii, Vanderwaltozyma polyspora, Tetrapisispora blattae, Saccharomyces kudriavzevii, Saccharomyces mikatae, Saccharomyces paradoxus, Saccharomyces cerevisiae*.

**Supplementary Figure 2.**
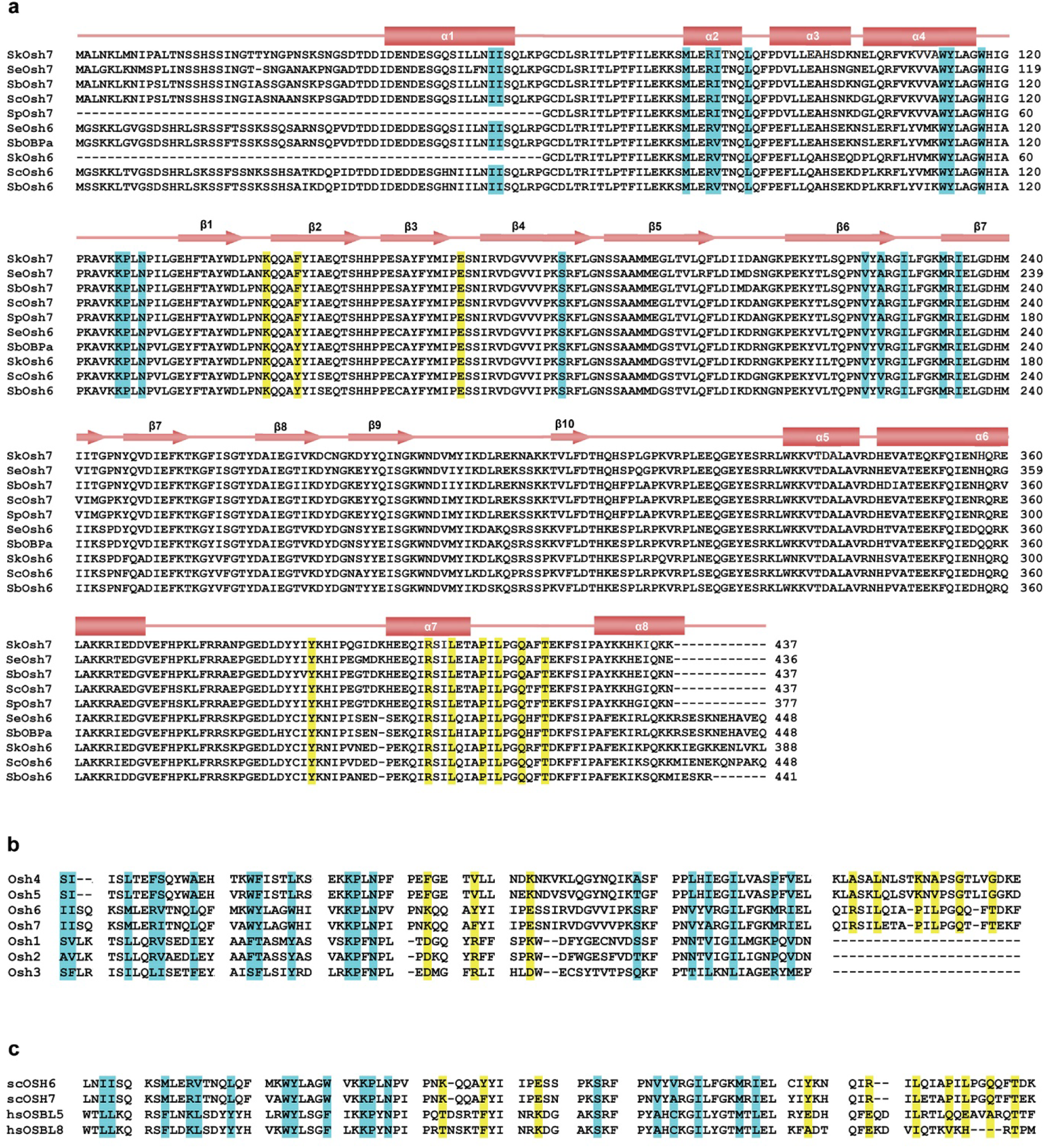
Sequence alignment of ORP proteins. **a**, Sequence alignment of Osh6 and Osh7 from five Saccharomyces species (Sp, *S. pastorianus*; Sk, *S. kudriavzeii*; Sc, *S. cerevisiae*; Se*, S. eubayanus*; Sb, *S. paradoxus*). The lipid (cyan) and Ist2 (yellow) binding sites are conserved. **b**, Sequence alignment of the regions constituting the Ist2 and PS binding sites of Osh6 in the seven Osh paralogs of *S. cerevisiae*. Lipid (cyan) and Ist2 (yellow) binding sites of Osh6 and Osh7 are not conserved in other paralogs. **c**, Sequence alignment of the regions constituting the Ist2 and PS binding sites of the *S. cerevisiae* paralogs Osh6 and 7 and the human OSP paralogs OSBL5 (ORP5) and OSBL8 (ORP8). Whereas there is strong conservation of the lipid binding site (cyan), the residues of the Ist2 binding site (yellow) are not conserved.

**Supplementary Figure 3.**
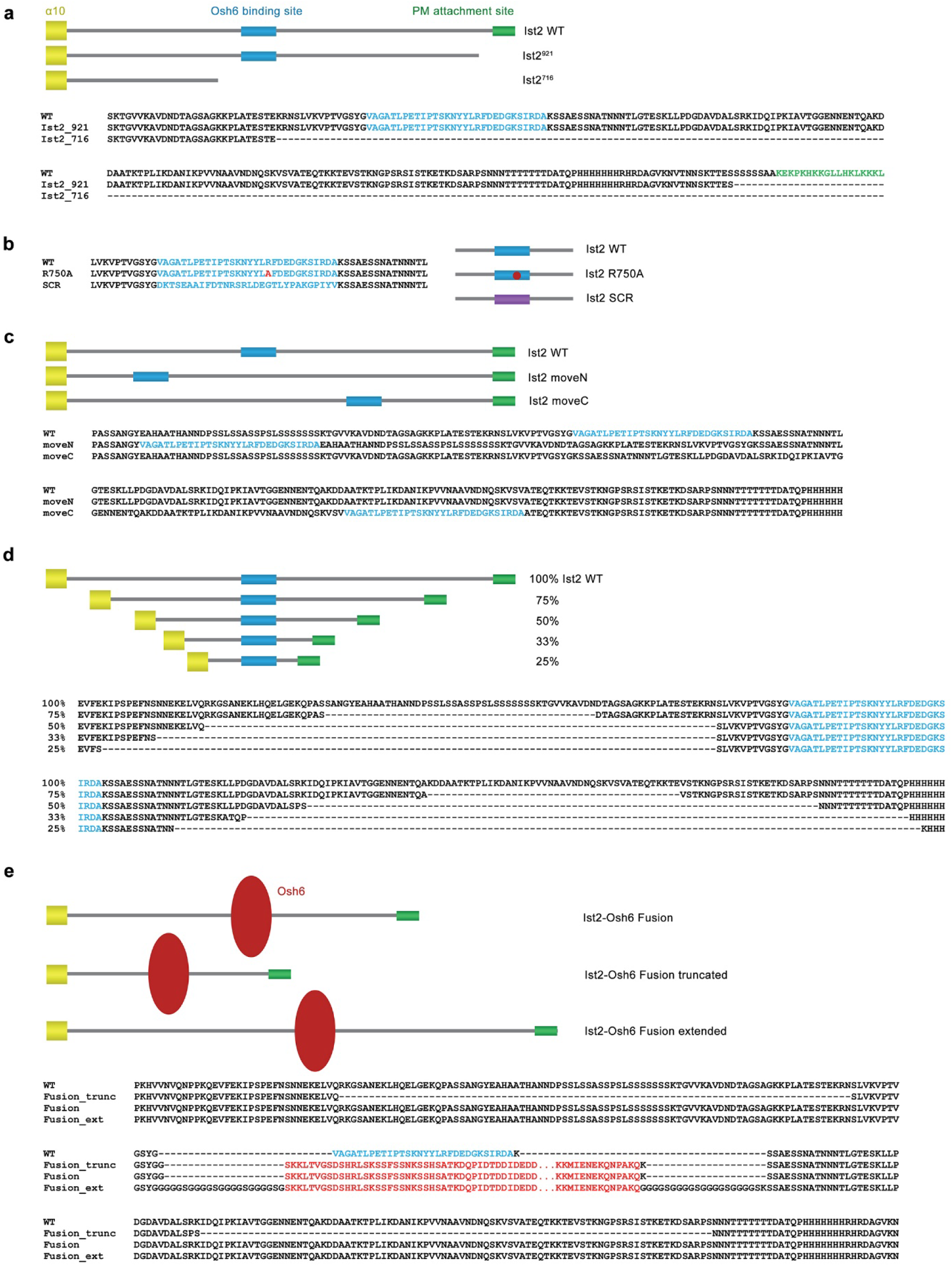
Yeast Complementation Assay constructs. **a,** Ist2 constructs with shortened C-terminus used to obtain data displayed in Fig. 5a. **b**, Ist2 constructs with mutated Osh6 binding site used to obtain data displayed in Fig. 5b. **c**, Ist2 constructs with displaced Osh6 binding site used to obtain data displayed in Fig. 5c. **d**, Ist2 constructs with truncated Ist2 linker used to obtain data displayed in Fig. 5d. **e**, Ist2 constructs with fused Osh6 used to generate data displayed in Fig. 5e. WT refers to full-length Ist2. Scheme of constructs is shown above. Residues of the Osh6 binding site are colored blue. Residue in red in (**b**) refers to location of point mutation, residues in red in (**e**) to Osh6 sequences.

